# Niche signalling regulates eIF3d1 phosphorylation to promote distinct modes of translation initiation in stem and differentiating cells

**DOI:** 10.1101/2023.12.15.571284

**Authors:** Ruoxu Wang, Mykola Roiuk, Freya Storer, Aurelio A. Teleman, Marc Amoyel

## Abstract

Stem cells have the unique ability among adult cells to give rise to cells of different identities. To do so, they must change gene expression in response to environmental signals. Much work has focused on how transcription is regulated to achieve these changes, however in many cell types, transcripts and proteins correlate poorly, indicating that post-transcriptional regulation is important. To assess how translational control can influence stem cell fate, we use the Drosophila testis as a model. The testis niche secretes a ligand to activate the JAK/STAT pathway in two stem cell populations, germline stem cells (GSCs) and somatic cyst stem cells (CySCs). We find that global translation rates are high in CySCs and decrease during differentiation, and that JAK/STAT signalling regulates translation. To determine how translation was regulated, we knocked down translation initiation factors and found that the cap binding complex, eIF4F, is dispensable in differentiating cells, but is specifically required in CySCs for self-renewal, acting downstream of JAK/STAT activity. Moreover, we identify eIF3d1 as a key regulator of CySC fate, and show that its phosphorylation is critical to maintain CySC self-renewal. We further show that Casein Kinase II, which controls eIF3d1 phosphorylation, is sufficient to restore CySC function in the absence of JAK/STAT. We propose a model in which niche signals regulate a specific translation programme in which only some mRNAs are translated, through regulation of eIF3d phosphorylation. The mechanism we identify allows stem cells to switch between modes of translation, adding a layer of regulation on top of transcription and providing cells with the ability to rapidly change gene expression upon receiving external stimuli.

## Introduction

To maintain adult tissue homeostasis, stem cells must have the ability to respond to the needs of the tissue in a timely manner to produce differentiating offspring. Observations across tissues suggest that a pool of stem cells exists in a poised, or licensed state, enabling them to rapidly transition to a differentiated state upon signals from the environment (Nakagawa et al., 2021; Rompolas et al., 2016; Yuen et al., 2021). Thus, the ability to switch gene expression profiles at short notice is critical to maintaining tissue integrity. How this is achieved is still poorly understood. Much work has focused on how transcription is regulated in stem and differentiating cells by signals from the niche, or supportive environment that maintains stem cell self-renewal, leading to an increasing understanding of gene regulatory networks and transcriptional landscapes during differentiation (Almeida et al., 2021; Davidson et al., 2002; Enver et al., 2009; Wilson et al., 2010; Young, 2011). However, comparison of the transcriptome and translatome has revealed that protein expression is poorly correlated with transcription in many stem cells and their differentiated progeny (Baser et al., 2019; Ingolia et al., 2011; Lu et al., 2009; Schwanhausser et al., 2011; Spevak et al., 2020). Therefore, to fully understand how cell identity transitions occur, it is critical to study the post-transcriptional control of gene expression and its regulation by niche signals promoting self-renewal.

An important point of control of gene expression is the translation of mRNA. Two broad classes of mechanisms affecting mRNA translation in stem cells have been described: in the first, which is best characterised in the germline stem cells (GSC) of the Drosophila ovary, sequence-specific RNA-binding proteins sequester and inhibit translation of mRNAs encoding differentiation factors (Blatt et al., 2020; Slaidina and Lehmann, 2014). The second class involves more global effects on translation rates which change during differentiation; surprisingly, these changes in global translation have major roles in determining cell fate, and act by preferentially affecting the translation of specific mRNAs, although in ways that are still poorly characterised (Saba et al., 2021; Wang and Amoyel, 2022). Thus, different translational programmes are overlaid onto the transcriptional programmes of stem cells and their differentiating progeny, leading to a complex regulation of gene expression. Importantly, it remains unknown how niche signals, which direct the transcriptional programme of stem cell gene expression, impact global translation to promote stem cell-specific translational programmes.

To explore the relationship between niche signals and translational regulation, we use the Drosophila testis as a model, since the niche and its signalling are well-understood and can be manipulated with exquisite precision (Greenspan et al., 2015). In the Drosophila testis, the stem cell niche, called the hub, is a cluster of 10-12 somatic cells that supports the self-renewal of two different stem cell populations: germline stem cells (GSCs) and somatic cyst stem cells (CySCs) (Greenspan et al., 2015; Hardy et al., 1979). GSCs give rise to daughter cells known as gonialblasts, which divide with incomplete cytokinesis to form germline cysts that eventually produce spermatids, while CySCs give rise to post-mitotic cyst cells which envelop gonialblasts and are critical for the development of germ cells. CySC self-renewal is primarily regulated by the Janus kinase (JAK)/signal transducer and activator of transcription (STAT) pathway. Hub cells secrete the JAK/STAT ligand Unpaired (Upd), which leads to activation of the sole Drosophila STAT, Stat92E, in stem cells immediately adjacent to the hub (Kiger et al., 2001; Tulina and Matunis, 2001). Active Stat92E acts to promote transcription of several targets that encode important factors for CySC self-renewal, including *Zn-finger homeodomain 1* (*zfh1*), and Zfh1 is commonly used as a marker for CySCs (Inaba et al., 2011; Issigonis et al., 2009; Leatherman and Dinardo, 2008). As they differentiate, cyst cells express Eyes absent (Eya) (Fig. 1A) (Fabrizio et al., 2003). The relative contributions of transcriptional and post-transcriptional regulation to these changes in gene expression downstream of JAK/STAT activation have not yet been determined.

**Figure 1.**
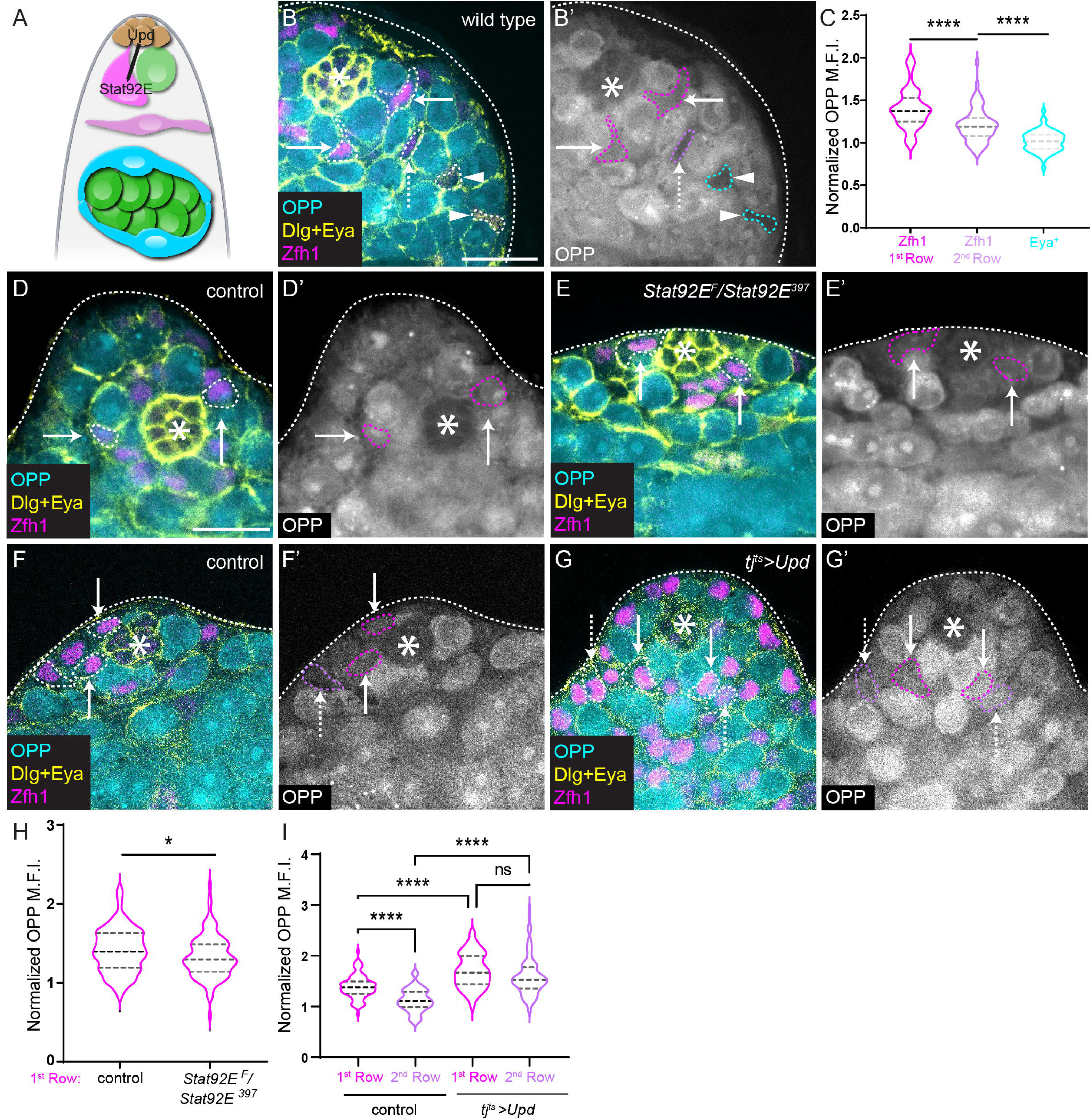
JAK/STAT signalling promotes high global translation rates in CySCs. (A) Schematic of a Drosophila testis. The hub (brown) produces the JAK/STAT ligand Unpaired (Upd) to activate Stat92E in surrounding stem cells. Two stem cell populations are supported by the hub, germline stem cells (green), and somatic cyst stem cells (CySCs) (magenta). CySCs are found in two rows and express Zfh1; they differentiate into cyst cells that express Eya (cyan). (B,D-G) OPP incorporation (cyan) was used to measure global translation rates in testes stained with antibodies against Dlg (yellow) to label the hub, Eya (yellow) to label fully differentiated cyst cells and Zfh1 (magenta) to label CySCs. Dashed lines outline CySCs adjacent to the hub (arrows), or in the 2^nd^ row away from the hub (dashed arrows) and fully differentiated cells (arrowheads). Asterisks mark the hub. (B,B’) A wild type testis showing highest OPP in CySCs adjacent to the hub and decreasing OPP levels with increasing distance from the hub. Scale bar: 15 µM. (C) Quantification of OPP incorporation in wild type CySCs, 2^nd^ row cyst cells and Eya^+^ cells, normalized to levels in the hub (N>80 cells from 10 testes, student’s t-test, **** P<0.0001.) (D-E’) A control (+/Stat92E^397^) testis (D,D’) and a testis from a temperature-sensitive Stat92e mutant (Stat92E^F^/Stat92E^397^) (E,E’) after 20h at the restrictive temperature. (F-G’) A control (tj^ts^ > +) testis (F,F’) and a testis with Upd over-expression (tj^ts^ > upd) (G,G’) after 20h at the restrictive temperature. (H) Quantification of OPP incorporation in 1^st^ row cyst cells (CySCs) normalized to levels in the hub. CySCs have significantly lower OPP levels in Stat92e mutant testes than in controls (N>100 cells from ≥8 testes, student’s t-test, * P<0.05). (I) Quantification of OPP incorporation in 1^st^ row and 2^nd^ row CySCs normalized to levels in the hub. Upd over-expression results in significantly higher OPP levels in both rows, and abolishes the difference between first and second row (N>80 cells from ≥7 testes, Šidák multiple comparisons test, **** P<0.0001, ns P=0.0946.)

Translation initiation is thought to be the rate-limiting step of protein synthesis and depends on several eukaryotic initiation factor (eIF) complexes (Jackson et al., 2010). Canonically, translation is initiated by binding of eIF4E to the 5’ m^7^G cap of the mRNA, where the eIF4F complex, composed of eIF4A, 4E and 4G subunits is assembled. eIF4F acts to recruit the eIF3 complex, a large multi-subunit initiation factor, which is bound to the ribosome and other eIFs, forming the 43S ribosome. This results in recruitment of the 43S to the 5’ end of the mRNA, where it begins to scan for a translation start site. There is increasing evidence, however, that eIFs are not merely scaffolds but that they are regulated and can selectively drive translation of specific transcripts. Non-canonical translation can occur upon sequestration of eIF4E by 4E-Binding Protein (4E-BP), or expression of a variant eIF4G, eIF4G2, also known as Death-Associated Protein 5 (DAP5), which lacks the eIF4E binding domain. In many cases, cap-independent translation is thought to occur through internal ribosome entry sites (IRES). Of note, the eIF3 subunit, eIF3d, can bind the cap in the absence of eIF4F and promote translation of mRNAs depending on the structure of the 5’ untranslated region (Lee et al., 2016; Ma et al., 2023). Indeed, it is thought that eIF3d drives cap-dependent translation together with DAP5 (de la Parra et al., 2018; Volta et al., 2021). Cap-binding activity of eIF3d is dependent on phosphorylation by Casein Kinase 2 (CK2) (Lamper et al., 2020), although to date this regulation has only been demonstrated in culture and in response to stress, and it is unknown whether it occurs during homeostasis in tissues. Non-canonical translation has been shown to occur in stem cells or during differentiation (Saba et al., 2021; Wang and Amoyel, 2022), yet how translation modes can be regulated in response to niche signals to influence cell fate is still poorly understood.

Here, we examined global translation in somatic cells of the testis. We found that translation levels are higher in CySCs than differentiating cyst cells, and that JAK/STAT signalling maintains high translation rates. To understand how these rates were regulated, we knocked down translation initiation factors and found a specific role for the mRNA 5’ m^7^G cap-binding eIF4F complex in maintaining self-renewal downstream of JAK/STAT, while most subunits of other initiation complexes were required for differentiation. Notably eIF3d1 was also required for self-renewal, and provides a potential mechanism for regulation of translation initiation since eIF3d is sensitive to phosphorylation (Lamper et al., 2020). Indeed, we show that a phospho-mimetic eIF3d1 can rescue loss of self-renewal upon Casein Kinase II (CkII) knockdown, while phospho-dead eIF3d1 cannot. Moreover, CK2 modulates the ability of eIF3d to interact with the eIF4F complex in human cells, suggesting that phosphorylation of eIF3d is a switch between different modes of translation initiation. Moreover, CkII over-expression is sufficient to functionally rescue CyCSs in the absence of JAK/STAT signalling, placing it as a key effector of self-renewal signalling. In sum, our data suggest that CySCs have a specific translational programme, requiring activity of the cap-binding eIF4F complex and that this activity is regulated by phosphorylation of eIF3d1 by CkII. The mechanism we uncover here suggests a model in which stem cells, loaded with messenger RNAs encoding both self-renewal and differentiation factors, can make rapid fate choices by switching translational programme, ensuring specificity in expression downstream of signals from the niche.

## Results

### JAK/STAT activity maintains high levels of translation in CySCs

Studies in several stem cell lineages have revealed that global translation rates change during differentiation (Saba et al., 2021; Wang and Amoyel, 2022). To monitor translation rates in the Drosophila testis, we incubated dissected testes with O-propargyl-puromycin (OPP) and visualised its incorporation in different cell types. Cells can be identified using molecular markers and by their position relative to the hub: CySCs contact the hub directly, and move further away as they differentiate (Hardy et al., 1979). Zfh1 labels CySCs and their immediate descendants, which include a population of cells that are licensed but not committed to differentiate, while differentiated cells express Eya (Fig. 1A) (Fabrizio et al., 2003; Leatherman and Dinardo, 2008; Yuen et al., 2021). OPP levels were higher in Zfh1-positive CySCs adjacent to the hub (Fig. 1B, arrows, quantified in Fig. 1C), and appeared to decline as cyst cells progressed through differentiation such that Zfh1-positive cells away from the hub (“2^nd^ row”, Fig. 1B dashed arrow, Fig. 1C) had lower OPP levels than those adjacent to the hub (14% decrease, P<0.0001), and Eya-positive cells had lower levels still (Fig. 1B, arrowhead, Fig. 1C, 16% decrease relative to 2^nd^ row cells, P<0.0001).

Given that JAK/STAT signalling is active in CySCs adjacent to the hub (Fig. 1A, (Issigonis et al., 2009; Leatherman and Dinardo, 2008) and essential for their self-renewal, we wondered whether the sole Drosophila STAT, Stat92E, could be responsible for maintaining high levels of translation in these cells. Shifting adult flies carrying a temperature-sensitive *Stat92E* allelic combination, referred to as *Stat92E^ts^*, to the restrictive temperature of 29°C results in differentiation of both CySCs and GSCs; however after one day of temperature shift, CySCs are still present, while GSCs begin to detach from the hub (Leatherman and Dinardo, 2010; Yuen et al., 2021), allowing us to address the role of JAK/STAT in regulating translation in CySCs. As assessed by OPP incorporation, we observed that 20 hours after temperature shift, global translation levels were decreased in Zfh1-positive CySCs immediately adjacent to the hub (Fig. 1D,E,H, 6% decrease compared to control, P<0.05), indicating that changes in translation occur following inactivation of Stat92E but prior to cell fate changes. Conversely, since Zfh1-positive cells exhibit different rates of global translation depending on their position relative to the hub, which is the source of the JAK/STAT ligand Upd, we asked whether providing ectopic Upd would increase translation rates away from the hub. We used the cyst lineage driver *traffic jam (tj)-Gal4*, together with a temperature-sensitive Gal80 (together referred to as *tj^ts^-Gal4*) to control the timing of expression and restrict it to adult stages. Indeed, over-expression of Upd resulted in higher levels of OPP incorporation in Zfh1-positive cells, both those contacting the hub and cells in the second row from the hub (22% and 42% increase, respectively, P<0.0001 for both relative to control, Fig. 1F,G,I). OPP levels in 2^nd^ row cells were indistinguishable from those in cells that contacted the hub (Fig. 1I). Thus, increasing JAK/STAT activity elevates translation rates in Zfh1-positive CySCs.

Altogether, our results indicate that global translation rates decrease as CySCs move away from the hub, and that the main signalling pathway mediating self-renewal in CySCs, JAK/STAT, maintains high levels of translation in CySCs. Thus, we sought to determine whether increased translation was important for the self-renewal of CySCs.

### Distinct requirements for translation initiation factors in CySCs and differentiating cells

To test whether high translation rates were necessary to maintain CySC identity, we asked how CySCs were affected if translation was impaired. Initiation is the most highly regulated stage of mRNA translation and involves the assembly of several eukaryotic initiation factor (eIF) complexes which recruit and activate ribosomes to scan the mRNA for the start codon (Fig. 2A) (Jackson et al., 2010). Thus, by disrupting translation initiation, we expected to reduce global translation rates, and if high translation was required for CySC self-renewal, these manipulations should result in loss of CySCs. We used *tj^ts^-Gal4* to knock down subunits of each eIF in somatic cells of the testis. Results of this screen are summarised in Fig. 2B and shown in full in Table S1.

**Figure 2.**
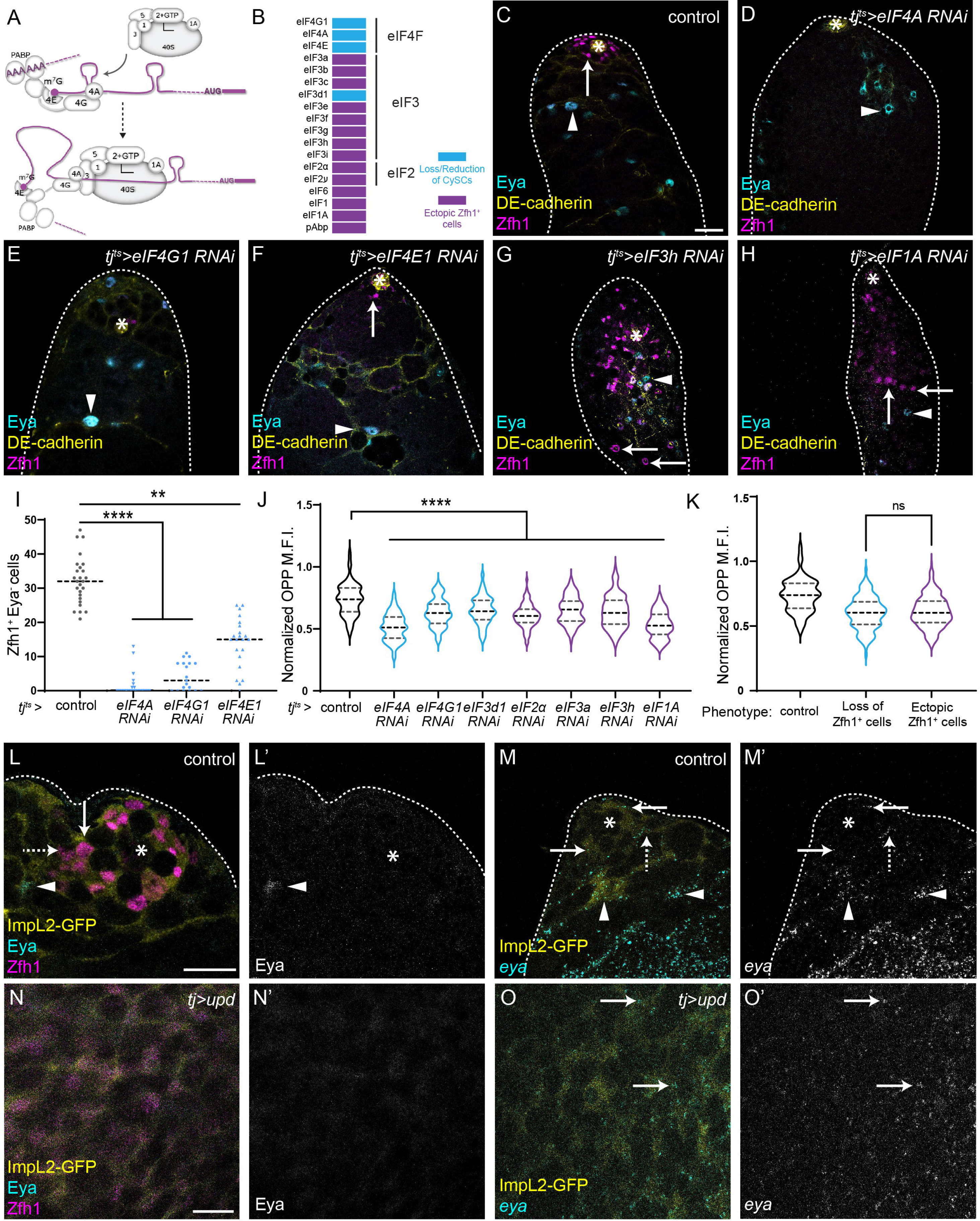
Knockdown reveals different requirements for translation initiation factors in CySCs and differentiating cyst cells. (A) Schematic of translation initiation. The eIF4F complex, composed of eIF4A, 4E and 4G, binds the 5’ m^7^G cap and interacts with the 43S pre-initiation complex, which consists of the multi-subunit eIF3 complex, the eIF2-initiator Methionine tRNA-GFT complex, and eIF1, 1A and 5, together with the small 40S ribosome subunit. Interactions between eIF4F and eIF3 result in the ribosome being brought to the 5’ end of the mRNA and scanning for the start codon. (B) Summary of the phenotypes observed upon knocking down eIFs. Knockdowns resulted either in complete loss or decrease of Zfh1-expressing cells (magenta), or ectopic presence of Zfh1-positive cells away from the hub (cyan). These phenotypes mostly clustered by complex, with the exception of eIF3d1. See Table S1 for full details. (C-H) Representative results from the initiation factor knockdown screen. A control (tj^ts^ > +) testis (C), and testes in which the indicated eIF was knocked down (D-H). DE-cadherin (yellow) labels the hub. Zfh1 (magenta) labels CySCs and Eya (cyan) labels differentiated cyst cells. Asterisks indicate the hub. Arrows indicate CySCs and arrowheads indicate differentiated cells. Scale bar: 15 µM. (I) Quantification of the number of Zfh1^+^ Eya^-^ cells upon knockdown of eIF4F complex subunits. (N≥19 testes, Kruskal-Wallis test, **** P<0.0001, ** P<0.01.) (J) Quantification of OPP incorporation in CySCs adjacent to the hub in each of the indicated genotypes, normalized to levels in GSCs. Knockdowns resulting in CySC loss are shown in magenta, those resulting in ectopic CySCs are shown in cyan, all lead to decreased OPP incorporation (N>85 cells from ≥8 testes, Šidák multiple comparisons test, **** P<0.0001.) (K) Grouped analysis of OPP levels in CySCs based on RNAi phenotype. No significant difference was observed between knockdowns resulting in CySC loss (magenta) or ectopic CySCs (cyan) (N>100 cells from ≥8 testes, Šidák multiple comparisons test, ns P=0.6764.) (L-O) Comparison of Eya protein (cyan L,N, single channel L’,N’) and eya mRNA (cyan M,O, single channel M’,O’) localisation in control testes (L,M) or testes over-expressing Upd and containing stem cell tumours (N,O). ImpL2-GFP (yellow) was used to identify CySCs together with Zfh1 antibody in immunostainings (L,N). Transcripts were detected in CySCs in control and in Upd over-expressing testes where no Eya protein is visible. Asterisks indicate the hub. Arrows mark CySCs and arrowheads mark differentiated cells. Scale bar: 15 µM.

Remarkably, we identified two opposite phenotypes, which clustered by complex. In the first, we observed either complete loss or severe reduction of CySC numbers, as assessed by Zfh1 expression. Knockdown of all subunits of the cap-binding eIF4F complex (eIF4A, eIF4E1, eIF4G1) as well as one subunit of eIF3, eIF3d1, resulted in significant decrease or complete absence of Zfh1-positive cells (Fig. 2D-F, quantified in Fig. 2I and see below for eIF3d1). In contrast, all other knockdowns led to the presence of ectopic Zfh1-positive cells away from the hub (Fig. 2G,H, Table S1 and see below).

These observations suggest that the eIF4F complex is necessary to maintain CySCs. To verify these findings and to test whether these phenotypes were cell autonomous, we generated positively-marked clones mutant for *eIF4A*. Control clones were observed at 2 days post clone induction (dpci) as GFP-positive cells adjacent to the hub (Fig. S1A, arrow). By 7 dpci, control clones were readily observed adjacent to the niche, indicating that the marked cells had self-renewed, and Eya-positive clonal cells were observed farther from the hub (Fig. S1B, arrows and arrowhead, respectively), indicating that the clones gave rise to differentiating cyst cells. In contrast, *eIF4A* mutant CySCs were rarely recovered at 2 dpci, and never at 7 dpci (Fig. S1C-F). This was not due to different induction rates, as marked cells (including both CySCs and differentiating cyst cells) were recovered at similar rates in mutant and control (Fig. S1E). Instead, all *eIF4A* mutant cells observed were also Eya-positive at 7 dpci (Fig. S1D, arrowhead), indicating that they had differentiated. To rule out the possibility that we did not observe *eIF4A* mutant CySCs because they were unable to synthesise GFP due to defective translation, we also generated negatively-marked clones.

These clones showed similar results; control CySC clones lacking GFP were observed at the hub at 2 dpci and maintained at 7 dpci, while *eIF4A* mutant CySC clones lacking GFP were occasionally observed adjacent to the niche at 2 dpci and almost never at 7 dpci (Fig. S1G-J). Additionally, we generated negatively-marked *eIF4E1* mutant clones, and observed a similar defect in mutant CySC clone recovery (Fig. S1K-N), indicating that, like *eIF4A, eIF4E1* is required autonomously for CySC self-renewal. The lack of marked mutant CySCs at 7 dpci could be due to two possibilities, either CySCs lacking eIF4A or eIF4E1 died, or they differentiated into cyst cells. Since we observed Eya-positive mutant cyst cells, we favoured the second possibility. Nonetheless, we used the baculovirus caspase inhibitor P35 to block apoptosis together with knockdown of eIF4G1 or eIF4A. Blocking cell death did not increase the number of CySCs compared to knockdown alone (Fig. S1O), indicating that cell death is not the primary cause of the reduction in CySCs upon knockdown of eIF4F subunits. Finally, we asked whether cyst cells lacking eIF4A could differentiate normally. Cyst cells encapsulate germ cells and support their development; as cyst cells differentiate, they acquire a flattened morphology that extends and surrounds germ cell cysts (Fig. S1P). We observed that *eIF4A* mutant cyst cells had a similar morphology, and germ cells encapsulated by these mutant cells appeared to develop normally (Fig. S1Q). Thus, we conclude that the cap-binding eIF4F complex is required specifically for self-renewal of CySCs, and that CySCs lacking eIF4F activity differentiate into functional cyst cells.

In contrast to loss of CySCs observed upon eIF4F knockdown, knockdown of subunits of all the other translation initiation complexes resulted in the presence of ectopic Zfh1-positive cells away from the hub (Fig. 2B, Table S1, examples are shown in Fig. 2G,H). These testes often also contained Eya-positive cells, indicating that some differentiation did occur. Nonetheless, the presence of ectopic Zfh1-positive cells away from the hub suggested that CySCs continued to self-renew when these translation initiation factors were knocked down. To confirm this, we assessed proliferation using the S-phase marker 5-ethynyl-2’-deoxyuridine (EdU), since CySCs are the only proliferating somatic cells in the testis (Cheng et al., 2011; Gonczy and DiNardo, 1996; Hardy et al., 1979). In controls, somatic EdU incorporation is only observed in Zfh1-positive cells adjacent to the hub (Fig. S2A, arrows). When eIF2 or eIF3 complex subunits, or other initiation factors such as eIF1, eIF1A and eIF6 were knocked down, we observed EdU incorporation in Zfh1-positive cells many cell diameters distant from the hub (Fig. S2B-D, dashed arrows), indicating that the ectopic Zfh1-positive cells have stem-like properties. Thus, inhibiting translation initiation factors other than eIF4F results in ectopic CySC self-renewal away from the niche and defective, although not fully blocked, differentiation.

In sum, knocking down translation initiation factors produced two opposite phenotypes: either ectopic self-renewal when most initiation factors were knocked down, or a loss of CySC self-renewal when eIF4F was knocked down. There are two possibilities to explain how such different phenotypes could be obtained when translation is perturbed: either different fates are induced by different levels of global translation, or knocking down different initiation factors selectively affects the translation of specific mRNAs. Firstly, given that we observed higher levels of translation in CySCs than in cyst cells (Fig. 1B,C), it is possible that these manipulations disrupt translation rates differently, such that eIF4F or eIF3d1 knockdown results in lower translation than other knockdowns, with a consequent effect on fate, where low translation rates lead to differentiation and high translation rates to self-renewal. To test this, we examined OPP incorporation in several conditions. All knockdowns we examined led to decreased translation relative to control (Fig. 2J); importantly, knockdowns that led to ectopic Zfh1-positive cells and knockdowns that led to loss of Zfh1-positive cells decreased OPP incorporation to a similar extent (Fig. 2K). Thus, we conclude that it is not the global rate of translation that determines cell fate. The alternative possibility to explain opposite phenotypes upon initiation factor knockdown is that translation initiation factors can determine which mRNAs are translated; in other words, that there are different translational programmes in CySCs and in differentiating cyst cells, determined by the activity of different translation initiation factors. For this model, two assumptions must be true: 1) that mRNA transcripts encoding both CySC and cyst cell factors must be present in the same cells, but only translated when appropriate, and 2) that translation initiation factor activity is regulated by signals promoting cell fate decisions to enable a switch from one programme to another.

To test the first assumption, we examined transcript localisation for *eya*. Eya protein is present in differentiating cyst cells and is absent from cells adjacent to the hub (Fig. 1A, 2L) (Fabrizio et al., 2003). However, previous work showed that *eya* mRNA was detectable in sorted CySCs, and the *eya* enhancer drives expression in CySCs (Leatherman and Dinardo, 2008; Ma et al., 2014; Sainz de la Maza et al., 2022). We therefore used *in situ* hybridisation chain reaction to examine the distribution of *eya* transcripts. We used *ImpL2-GFP* to identify CySCs (Amoyel et al., 2016; Terry et al., 2006). As expected, we observed many punctae of *eya* signal in differentiating cyst cells (Fig. 2M arrowheads). Notably, *eya* transcripts were also detected in GFP-positive cells in the first and second rows from the hub (Fig. 2M arrows), despite the fact that Eya protein is not detected in cells adjacent to the hub (Fig. 2L arrows). To confirm this result, we generated tumours consisting of only CySCs and GSCs by over-expressing Upd (Leatherman and Dinardo, 2008). In these tumours, all somatic cells expressed ImpL2-GFP and Zfh1, while Eya protein was rarely, if ever detected (Fig. 2N), consistent with previous observations (Amoyel et al., 2013; Leatherman and Dinardo, 2008). In contrast, we consistently detected *eya* transcripts in somatic cells, marked by ImpL2-GFP (Fig. 2O). Therefore, at least in the case of *eya*, transcripts encoding differentiation factors are expressed in stem cells, while their encoded protein is not, consistent with the existence of a selective translational programme in CySCs. We next set out to test the second assumption, that translation initiation is regulated downstream of niche signals.

### eIF4F functions downstream of JAK/STAT in CySC self-renewal

Thus far, our results indicate that there are different requirements for translation initiation factors in CySCs and differentiating cyst cells, and that transcripts encoding differentiation factors are present in stem cells, supporting a model in which regulation of initiation factors determines which transcripts are translated. Moreover, we showed that high global translation rates in CySCs depend on JAK/STAT activity (Fig. 1). Since initiation is the rate-limiting step in translation, this suggests that the CySC self-renewal pathway JAK/STAT could act to regulate CySC-specific translation initiation. However, since both eIF4F and JAK/STAT loss-of-function result in loss of CySC self-renewal, another possibility is that eIF4F is required for the translation of JAK/STAT signal transduction components and that signalling is compromised when eIF4F subunits are knocked down. To rule out this possibility, we examined levels of Stat92E, which is stabilised only in cells where signalling is active (Flaherty et al., 2010). We generated clones marked by loss of GFP and measured Stat92E immunofluorescence in clones normalised to neighbouring non-clonal cells (Fig. 3A,B quantified in 3F). We did not observe any decrease in Stat92E levels in *eIF4A* null mutant CySCs compared to control CySCs, indicating that JAK/STAT signalling is functional in mutant cells and cannot account for the self-renewal defect of eIF4F-deficient CySCs.

**Figure 3.**
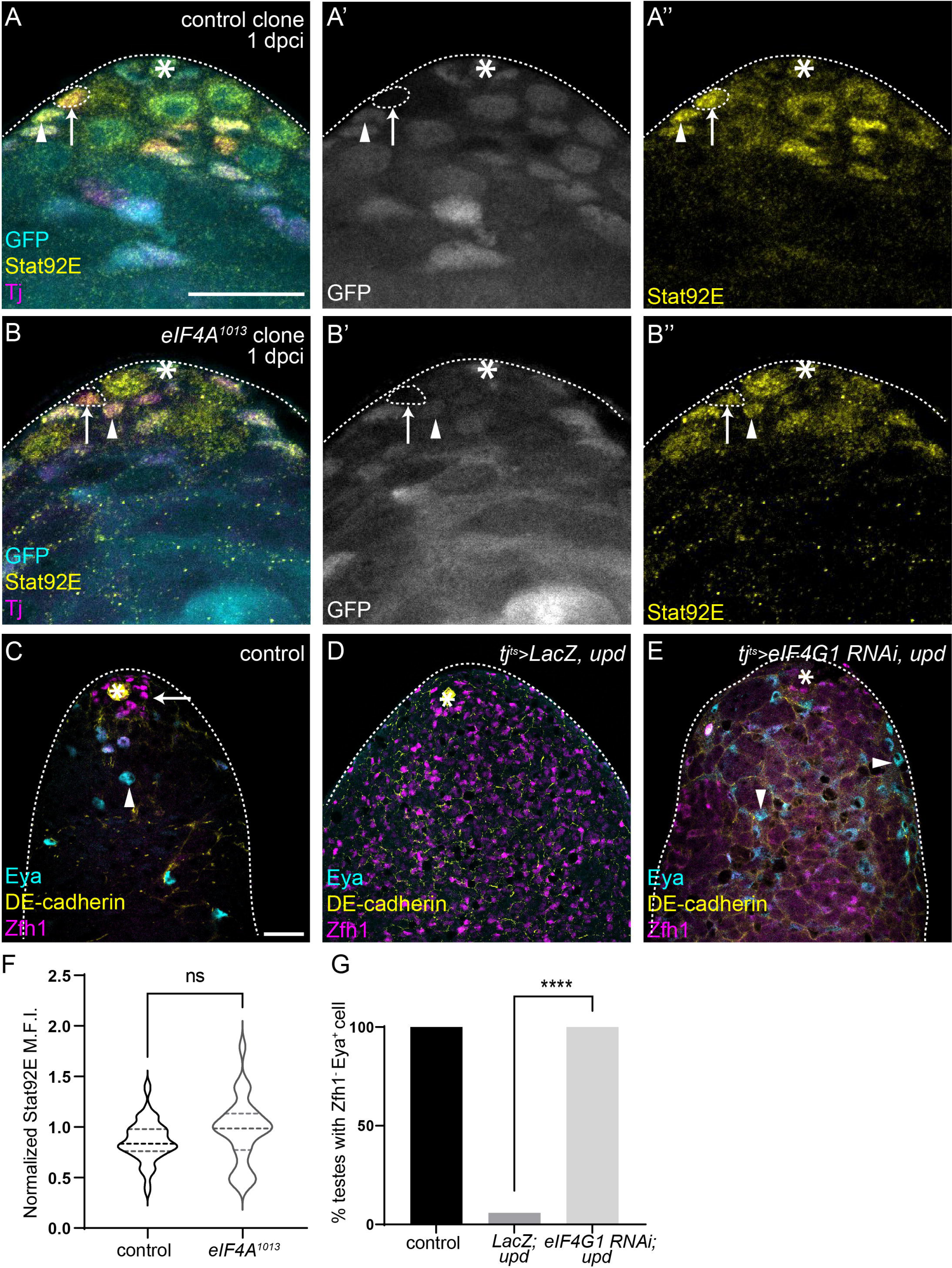
eIF4F is epistatic to JAK/STAT for CySC self-renewal. (A-B’’) Testes with control clones (A-A’’) and eIF4A^1013^ clones (B-B’’), 2 days post clone induction (dpci). Clones are marked by absence of GFP (cyan), and Tj (magenta) labels CySCs and early cyst cells. Dashed lines outline clonal CySCs (arrows) and arrowheads mark non-clonal CySCs. Asterisks indicate the hub. Scale bar: 15 µM. (C-E) A control (tj^ts^ > LacZ, +) testis (C), a testis in which Upd is over-expressed (tj^ts^ > LacZ, upd) (D) and a testis with concomitant Upd over-expression and eIF4G1 knockdown in CySCs (tj^ts^ > eIF4G1 RNAi, upd) (E). Upd over-expression results in stem cell tumours containing only Zfh1-positive cells and a lack of differentiated Eya-positive cells, while eIF4G1 knockdown rescues Eya-positive cells. DE-cadherin (yellow) labels the hub. Zfh1 (magenta) labels CySCs and Eya (cyan) labels differentiated cyst cells. Arrows mark CySCs and arrowheads mark differentiated cells. Asterisks indicate the hub. Scale bar: 15 µM. (F) Quantification of Stat92E levels in CySC clones (arrows in A-B’’) of the indicated genotype, normalized to levels in neighbouring non-clonal CySCs (arrowheads in A-B’’) (N≥28 cells from ≥11 testes, Student’s t-test, ns P=0.0943). (G) Percentage of testes with Eya-positive, Zfh1-negative differentiating cyst cells in the indicated genotypes (N≥15 testes, Chi-square test, **** P<0.0001).

Next, we sought to establish whether translation initiation through eIF4F is required downstream of JAK/STAT signalling for CySC self-renewal. Over-expression of the JAK/STAT ligand Upd in testes using *tj^ts^-Gal4*, together with LacZ as a titration control, resulted in tumours containing many Zfh1-positive CySC-like cells, and an almost complete absence of Eya-positive differentiated cyst cells (Fig. 3C,D). By contrast, knocking down eIF4G1 restored the presence of Eya-positive cells in all testes examined (Fig. 3E,G), indicating that eIF4F is required for self-renewal even in the presence of ectopic niche signals. We note that ectopic Zfh1-positive stem cells were still present and intermingled with Eya-positive cells in these testes, indicating an incomplete suppression of the JAK/STAT gain-of-function phenotype, and suggesting that translation is not the only downstream effector of JAK/STAT signalling. Nonetheless, given the known role of JAK/STAT in regulating transcription, it is striking that modulating translation alone was sufficient to alter cell fate when JAK/STAT was hyperactivated.

### eIF3d1 is required for CySC self-renewal

Our results suggest that eIF4F activity is modulated such that it is specifically required for translation in CySCs but not during differentiation, suggesting that there is a mechanism to selectively engage eIF4F in CySCs. One potential regulator of eIF4F activity is the eIF4E-binding protein (4E-BP) which binds eIF4E and prevents it from binding to the 5’ mRNA cap (Lin et al., 1994; Pause et al., 1994). Target of Rapamycin (Tor) inactivates 4E-BP by phosphorylation, allowing eIF4E to interact with mRNAs (Brunn et al., 1997; Gingras et al., 1999). However, we previously showed that 4E-BP is phosphorylated most highly in cells that leave the niche (Amoyel et al., 2016; Yuen et al., 2021). Therefore p4E-BP levels are high in cells that are likely to differentiate, which is inconsistent with the results described here in which eIF4F activity is specifically required for self-renewal. Thus, 4E-BP is unlikely to be the switch controlling eIF4F activity in CySCs.

While searching for an alternative mechanism that could be responsible, we noticed in our data above (Fig. 2B) that, in addition to eIF4F, only one other initiation factor was required for CySC maintenance: eIF3d1, the Drosophila homologue of eIF3d. This was notable for two reasons: firstly, eIF3d is one of the subunits that mediates the interaction between eIF4F and eIF3 (Brito Querido et al., 2020; Sun et al., 2011; Villa et al., 2013). Secondly, in cell culture, phosphorylation of eIF3d has been shown to act as a molecular switch for eIF3d function, suggesting a mechanism by which eIF3d could alter translation in response to environmental changes (Lamper et al., 2020). Thus, to determine whether eIF3d1 could be regulated to control translation and maintain CySC self-renewal, we characterised the *eif3d*1 loss-of-function phenotype. Knockdown of eIF3d1 in adult testes resulted in loss of Zfh1-positive cells (Fig. 4A,B), such that many testes were entirely devoid of CySCs (Fig. 4C, P<0.0001). To ensure this phenotype was autonomous to CySCs, we generated mutant clones homozygous for two independent alleles of *eif3d1*. Control clones were observed adjacent to the hub at 2 dpci (Fig. 4D) and maintained at 7 dpci, when they consisted of both Zfh1-positive cells and Eya-positive cells (Fig. 4E, arrow and arrowhead, respectively). *eif3d1* mutant clones were induced at similar rates to control (Fig. 4F), but were rarely observed adjacent to the hub (Fig. 4G). By 7 dpci, very few mutant CySC clones were recovered, and mutant cells were Eya-positive (Fig. 4H, arrowhead), indicating that they had differentiated. Expression of the caspase inhibitor P35 could not restore CySC numbers upon *eif3d1* knockdown (Fig. S1O). Altogether the clonal and knockdown data suggest that eIF3d1 is required autonomously in CySCs for self-renewal and that CySCs lacking eIF3d1 were lost from the niche due to differentiation, not cell death.

**Figure 4.**
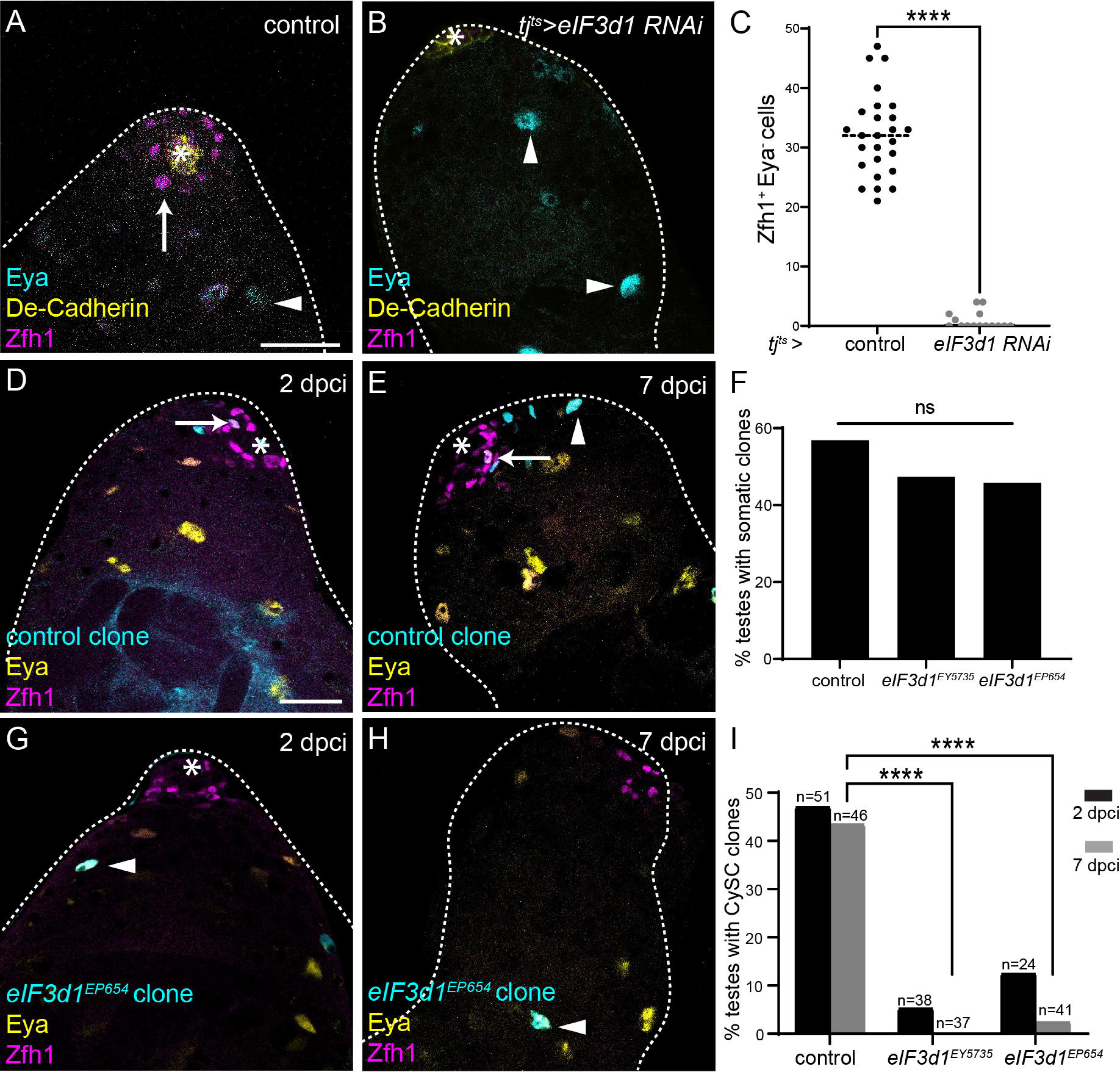
eIF3d1 maintains CySC self-renewal. (A-B) A control (tj^ts^ > +) testis (A) and testis in which eIF3d1 was knocked down somatically (tj^ts^ > eIF3d1 RNAi) (B). DE-cadherin (yellow) labels the hub. Zfh1 (magenta) labels CySCs and Eya (cyan) labels differentiated cyst cells. Arrows mark CySCs and arrowheads mark differentiated cells. Asterisks indicate the hub. Scale bar: 15 µM. (C) Number of Zfh1^+^ Eya^-^ cells in testes of the indicated genotype (N>=16 testes, Mann-Whitney test, **** P<0.0001). (D-E) Testes with control clones at 2dpci (D) and 7dpci (E), marked by GFP expression (cyan). Zfh1 (magenta) labels CySCs and Eya (yellow) labels differentiated cells. Arrows mark clonal CySCs and arrowheads mark clonal differentiated cells. Asterisks indicate the hub. Scale bar: 15 µM. (F) Percentage of testes with clones in either CySCs or differentiated cyst cells at 2 dpci (N≥24 testes, Chi-square test, ns P=0.5616). (G-H) Testes with eIF3d1^EP654^ clones at 2dpci (G) and 7dpci (H) marked by GFP expression (cyan). Zfh1 (magenta) labels CySCs and Eya (yellow) labels differentiated cells. Arrowheads mark clonal differentiated cells; note that mutant CySCs are not observed. Asterisks indicate the hub. Scale bar: 15 µM. (I) Percentage of testes with marked CySC clones at 2 dpci and 7 dpci. In controls, clones are induced and maintained at 7 dpci, while for both alleles of eIF3d1, few mutant CySC clones are recovered (N≥24 testes, Chi-square test, **** P<0.0001).

### Phosphorylation of eIF3d1 by CkII maintains CySC self-renewal

Previous work has shown that phosphorylation of eIF3d can promote translation in two different modes: as part of the canonical eIF3 complex, which binds eIF4F and promotes eIF4E-dependent translation, or alternatively as a cap-binding protein that promotes translation when eIF4E is inactivated (Lee et al., 2016). Phosphorylation of eIF3d has been shown to regulate its cap-binding activity: unphosphorylated eIF3d binds the cap while phosphorylated eIF3d does not (Lamper et al., 2020). In mammalian cells, Casein Kinase 2 (CK2, CkII in Drosophila) is the kinase responsible for phosphorylating eIF3d (Lamper et al., 2020). These phosphorylation sites are predicted to be conserved in Drosophila (Fig. S3A).

We therefore tested whether the Drosophila CK2 homologue, CkII, composed of α and β subunits, was required for eIF3d1 function in CySCs. Knockdown of either *CkIIα* or *CkIIβ* using two independent lines with *tj^ts^-Gal4* led to a dramatic loss of Zfh1-positive CySCs, with many testes entirely devoid of CySCs although Eya-positive cyst cells were observed (Fig. 5A-D). Thus, CkII is required for CySC self-renewal, similarly to eIF3d1. To determine whether CkII affected translation in CySCs, we measured OPP incorporation in CySCs after 1 day of *CkIIβ* knockdown, when CySCs were still present at the hub. The rate of global translation was significantly reduced in CySCs in which *CkIIβ* was knocked down compared to controls (Fig. 5E, 14% reduction compared to control, P<0.0001), suggesting that decreases in translation occur upon CkII knockdown prior to loss of CySCs.

**Figure 5.**
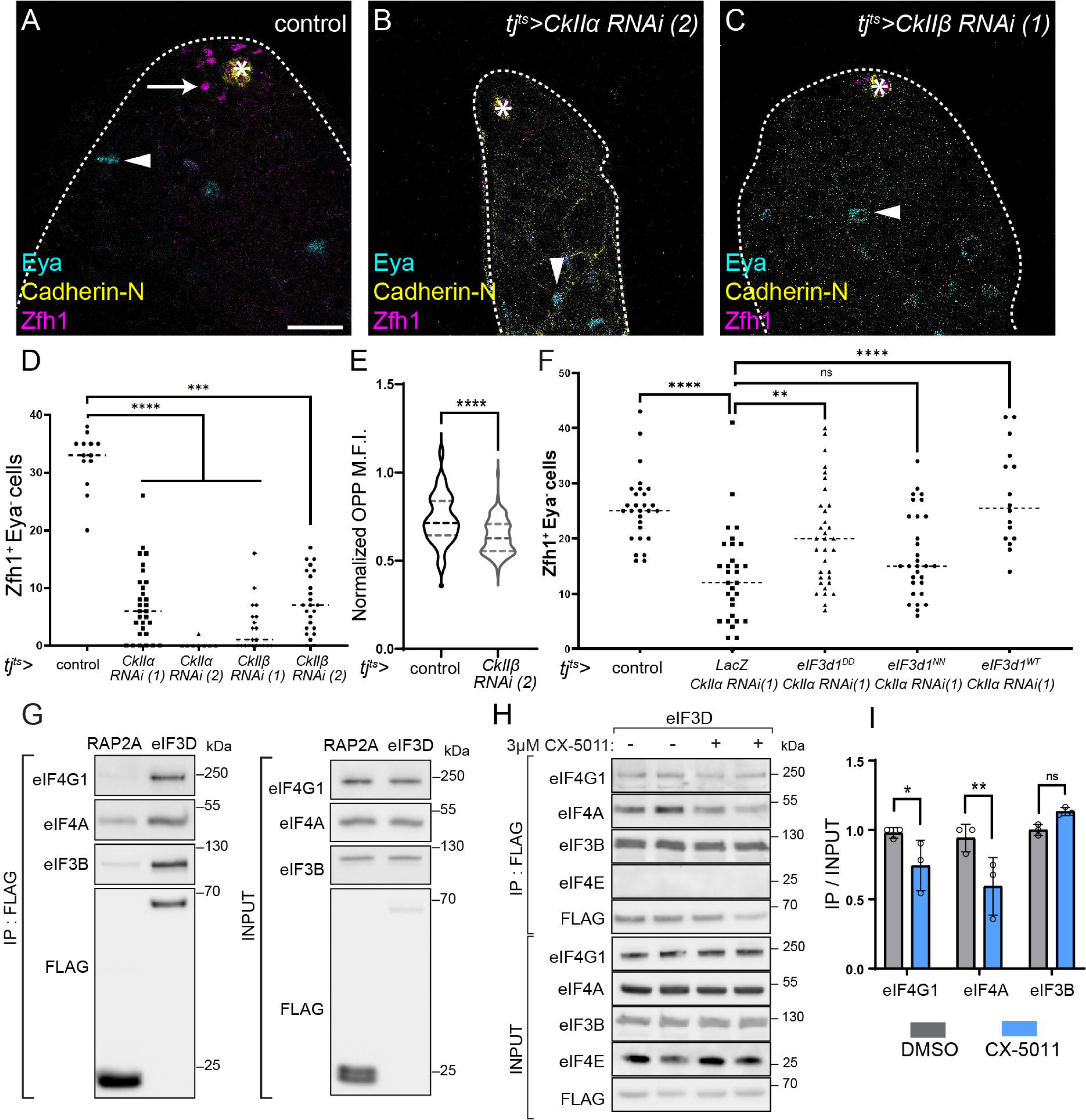
CkII promotes the interaction between eIF3d1 and eIF4F to maintain CySC self-renewal. (A-C) A control (tj^ts^ > LacZ, +) testis (A), and testes in which CkIIα (B) or CkIIβ was knocked down (C). Cadherin-N (yellow) labels the hub. Zfh1 (magenta) labels CySCs and Eya (cyan) labels differentiated cyst cells. Arrows mark CySCs and arrowheads mark differentiated cells. Asterisks indicate the hub. Scale bar: 15 µM. (D) Number of Zfh1^+^ Eya^-^ CySCs in testes of the indicated genotypes. Two independent RNAi lines were used to knock down both CkIIα and CkIIβ, leading to significant loss of CySCs (N≥7 testes, Kruskal-Wallis test, **** P<0.0001, *** P<0.001). (E) Quantification of OPP incorporation in CySCs adjacent to the hub in control or CkIIβ knockdown, normalized to levels in GSCs (N>70 cells from ≥8 testes, student’s t-test, **** P<0.0001). (F) Number of Zfh1^+^ Eya^-^ CySCs in testes in which CkIIα was knocked down, together with over-expression of phospho-mimetic (eIF3d1^DD^), phospho-dead (eIF3d1^NN^) or wild type eIF3d1 (eIF3d1^WT^). Phospho-mimetic and wild type eIF3d1 expression significantly rescue CySC numbers while phospho-dead eIF3d1 does not (N≥16 testes, Kruskal-Wallis test, **** P<0.0001, ** P<0.01, ns P=0.2892). (G) Western blot from lysates of control (RAP2A-FLAG) or eIF3d-FLAG-transfected cells, blotted with antibodies against eIF4G1, eIF4A, eIF3B and FLAG, either in total lysate (input, right) or after immunoprecipitation with an anti-FLAG antibody (IP, left). eIF3d pull down enriched for other initiation factors. (H) Western blot from lysates of eIF3d-FLAG-transfected cells, blotted with antibodies against eIF4G1, eIF4A, eIF3B and FLAG, either in total lysate (input, right) or after immunoprecipitation with an anti-FLAG antibody (IP, left). The two replicates on the right of each blot were incubated with the CK2 inhibitor CX-5011; in this condition, less eIF4G1 and eIF4A was immunoprecipitated with eIF3d-FLAG. (I) Quantification of immunoprecipitation blots showing that less eIF4G1 or eIF4A is co-immunoprecipitated with eIF3d-FLAG upon incubation with CX-5011. (N=3 replicates, student’s t-test, * P<0.037, ** P<0.005).

Next, we asked whether phosphorylation of eIF3d1 was required downstream of CkII for CySC self-renewal. We reasoned that if the role of CkII was to phosphorylate eIF3d1, then providing an exogenous source of phosphorylated eIF3d1 should rescue the loss of CySCs caused by CkII knockdown. We generated UAS-eIF3d1 constructs in which the predicted CkII target serines were mutated to produce a phospho-mimetic form of eIF3d1 (UAS-eIF3d1^DD^) or an unphosphorylatable form (UAS-eIF3d1^NN^) and over-expressed wild-type, phospho-mimetic or phospho-dead eIF3d1 while knocking down CkII. In testes in which *CkIIα* was knocked down alone, along with a suitable titration control, CySC numbers were reduced compared to control (P<0.0001, Fig. 5F). Expression of phospho-dead eIF3d1 had no effect, with CySC numbers similar to the knockdown control (UAS-eIF3d1^NN^, P=0.29, Fig. 5F). By contrast, expression of phospho-mimetic eIF3d1 led to a significant rescue of CySCs (UAS-eIF3d1^DD^, P<0.01, Fig. 5F). Unexpectedly, expression of wild type eIF3d1 also led to a significant rescue (UAS-eIF3d1^WT^, P<0.0001, Fig. 5F), suggesting that other kinases may regulate eIF3d1 phosphorylation non-redundantly with CkII, but that eIF3d1 levels are limiting in this context. Consistent with a requirement for eIF3d1 phosphorylation in CySC self-renewal, expression of wild type or phospho-mimetic eIF3d1 had no effect on CySC numbers (Fig. S3B); however, expression of phospho-dead eIF3d1 resulted in a 15% decrease in the number of CySCs (P<0.05), suggesting that this construct acts as a dominant-negative. eIF3d can bind the mRNA 5’ cap directly to drive translation of specific transcripts (Lee et al., 2015; Lee et al., 2016). We expressed a mutant form of eIF3d1 lacking the cap-binding domain, eIF3d1^helix11^, which can act as a dominant-negative construct when eIF3d1 cap-binding function is required (Lee et al., 2016; Rode et al., 2018). However, expression of this construct did not affect CySC numbers (Fig. S3C).

Our data indicate that eIF4F is required for CySC self-renewal, suggesting that canonical cap-dependent translation occurs in CySCs. Loss-of-function of CkII or of eIF3d1 phenocopy eIF4F loss suggesting that it is not the cap-binding activity of eIF3d1 that is required in CySCs. These observations suggest a new model for eIF3d1 regulation by CkII, where phosphorylation of eIF3d1 is necessary to maintain the interaction between eIF4F and eIF3d1, in addition to the previously-described role for this modification in modulating the cap-binding ability of eIF3d1 (Lamper et al., 2020). This would result in eIF3d1 acting as a switch to mediate recruitment of the 43S ribosome to mRNAs bound to eIF4F and would be consistent with loss-of-function of CkII, eIF3d1 and eIF4F subunits having similar phenotypes.

To test this hypothesis, we asked whether CK2 activity influenced the ability of eIF3d to interact with eIF4F using human cells in culture. We immunoprecipitated FLAG-tagged eIF3d and found, as expected, that it could co-immunoprecipitate eIF4A and eIF4G1 (Fig. 5F), consistent with structural models and biochemical data placing eIF3d at the interface between the eIF3 and eIF4F complexes (Brito Querido et al., 2020; Sun et al., 2011; Villa et al., 2013). Next we treated cells with the CK2 inhibitor CX-5011, and repeated the pull-downs. We confirmed that CX-5011 inhibited CK2 by blotting total cell lysates with a pan-phospho-CK2 substrate antibody (Fig. S3D). In these cells, CK2 inhibition resulted in a significant decrease in the ability of FLAG-tagged eIF3d to co-precipitate eIF4A and eIF4G1, (37% and 24% decrease IP relative to input, P<0.005 and P<0.037, respectively, Fig. 5G,H). Notably, the interaction between eIF3d and eIF3b was unaffected by CK2 inhibition (Fig. 5G,H). Thus, CK2 is required for maximal association of eIF3d with eIF4F, but eIF3d association with the eIF3 complex appears independent of regulation by CK2. Altogether, our data suggest a new model for translational regulation by eIF3d1 in CySCs: CkII phosphorylates eIF3d1 to promote its interaction with the eIF4F complex.

### CkII is epistatic to JAK/STAT in CySCs

Finally, since the JAK/STAT pathway is the main regulator of self-renewal and we showed above that its activity maintains high translation in CySCs, we asked whether JAK/STAT could regulate translation through CkII. First, we tested whether CkII was required genetically downstream of JAK/STAT to maintain CySC self-renewal. We over-expressed the JAK/STAT pathway ligand Upd using *tj^ts^-Gal4*, giving rise to tumours composed of CySCs expressing Zfh1 and devoid of Eya-expressing cyst cells (Fig. 6A). When *CkIIα* was knocked down in these testes, only Eya-positive cells were detected, and no Zfh1-expressing CySCs were present (Fig. 6B,C), although in rare cases double positive differentiating cyst cells were observed (Fig. 6C). Thus, CkII is required downstream of JAK/STAT for CySC self-renewal. We ruled out the possibility that this requirement was because CkII affected the transduction of JAK/STAT signalling by monitoring levels of Stat92E and saw no difference between control and CkII knockdown CySCs (Fig. S4).

**Figure 6.**
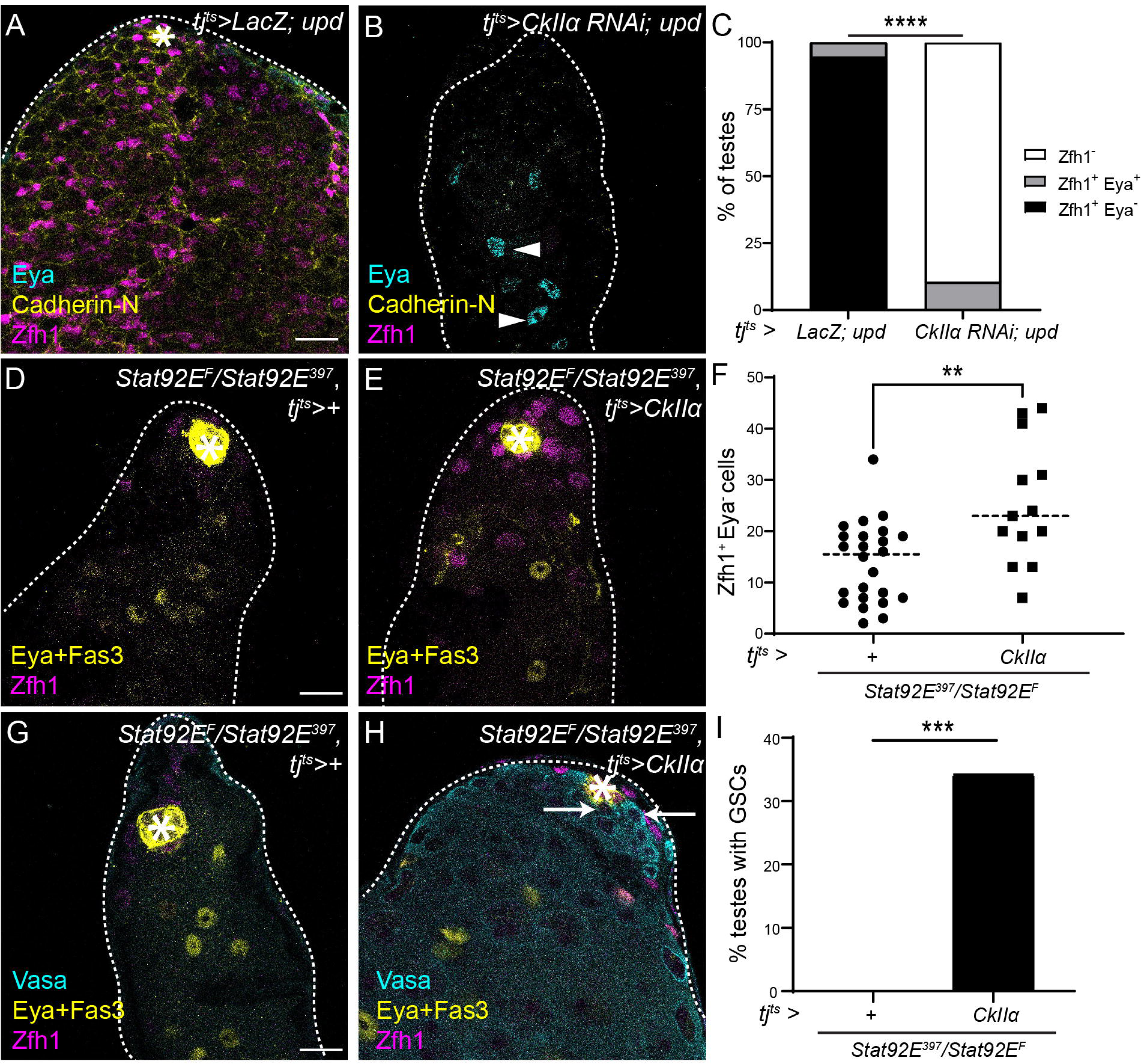
CkII mediates both autonomous and non-autonomous functions of JAK/STAT signalling in CySCs. (A,B) A testis in which JAK/STAT is hyperactivated (tj^ts^ > LacZ, upd) contains only CySCs and is devoid of differentiating cyst cells (A) while a testis in which CkIIα is knocked down in CySCs concomitantly with JAK/STAT hyperactivation (tj^ts^ > CkIIα RNAi, upd) contains only Eya-positive differentiated cyst cells (B). Cadherin-N (yellow) labels the hub. Zfh1 (magenta) labels CySCs and Eya (cyan) labels differentiated cyst cells. Arrows mark CySCs and arrowheads mark differentiated cells. Asterisks indicate the hub. Scale bar: 15 µM. (C) Percentage of testes containing only CySCs (Zfh1^+^ Eya^-^, black), both CySCs and cyst cells (Zfh1^+^ Eya^+^, grey) and no CySCs (Zfh1^-^, white) in the indicated genotypes (N≥18 testes, Chi-square test, **** P<0.0001). (D,E) A testis from a temperature-sensitive Stat92e mutant (tj^ts^>+; Stat92E^F^/Stat92E^397^) (D) and a testis from a temperature-sensitive Stat92e mutant expressing CkIIα (tj^ts^ > CkIIα, Stat92E^F^/Stat92E^397^) (E) after 10 days at the restrictive temperature. CkIIα expression results in increased numbers of CySCs around the hub. Fas3 (yellow) labels the hub, Zfh1 (magenta) labels CySCs and Eya (yellow) labels differentiated cyst cells. Asterisks indicate the hub. Scale bar: 15 µM. (F) Number of Zfh1^+^ Eya^-^ CySCs in testes of the indicated genotype (N≥13 testes, Mann-Whitney test, ** P<0.01). (G,H) Germ cells (Vasa, cyan) are lost in a testis from a temperature-sensitive Stat92e mutant (tj^ts^>+; Stat92E^F^/Stat92E^397^) (G) but are clearly visible in a testis from a temperature-sensitive Stat92e mutant expressing CkIIα (tj^ts^ > CkIIα, Stat92E^F^/Stat92E^397^) (H) after 10 days at the restrictive temperature. Fas3 (yellow) labels the hub. Zfh1 (magenta) labels CySCs and Eya (yellow) labels differentiated cyst cells. Arrows indicate GSCs, identified as individual round Vasa-positive cells adjacent to the hub. Scale bar: 15 µM. (I) Percentage of testes containing any GSCs in temperature-sensitive Stat92e mutants alone or expressing CkIIα (N≥30 testes, Chi-square test, *** P<0.001).

Next, we tested whether expressing the catalytic subunit of CkII, CkIIα, was sufficient to maintain CySC function in the absence of JAK/STAT signalling. We used the *Stat92E^ts^* mutant and observed after 10 days at the restrictive temperature that there were few CySCs around the hub, albeit in this genetic background, CySCs were not completely lost (Fig. 6D). Expression of the catalytic subunit of CkII, CkIIα, with *tj^ts^-Gal4* led to an increase in CySC numbers (Fig. 6E,F, P<0.01), suggesting that CkII can sustain CySC self-renewal in the absence of JAK/STAT signalling. Strikingly, we noticed that testes from *Stat92E^ts^* mutants in which CkIIα was over-expressed appeared to contain many more cells. In *Stat92E^ts^* mutants, GSCs are lost as well as CySCs, but previous work showed that restoring Stat92E function only in CySCs was sufficient to maintain both stem cell populations (Leatherman and Dinardo, 2010). Therefore, we examined germ cell presence and morphology using an antibody against Vasa in these testes. In agreement with previous findings, *Stat92E^ts^* mutants shifted to the restrictive temperature for 10 days lacked GSCs, identified as individual round cells in contact with the hub, and were often devoid of spermatogonia entirely (Fig. 6G-I) (Leatherman and Dinardo, 2010). In contrast, testes in which CkIIα was expressed often contained GSCs and appeared full of spermatogonia (Fig. 6H,I). Thus, expression of CkIIα in CySCs was sufficient to compensate for both autonomous functions of JAK/STAT in CySC self-renewal and non-autonomous functions in maintaining GSCs.

## Discussion

Despite extensive evidence that translation is regulated in stem cells to influence fate decisions, there is little understanding to date of how this regulation is achieved in response to signals from the stem cell niche. Here, using the somatic CySCs of the Drosophila testis as a model, we uncover such a regulation of translation by niche signals, and we propose that such post-transcriptional mechanism achieves two goals: firstly, it allows stem cells to selectively translate (or not translate) mRNAs important for determining cell identity, ensuring that transcriptional noise does not result in aberrant protein expression; secondly, by enabling cells to switch between translational programmes and by having pre-existing expression of mRNAs encoding differentiation factors, this regulation allows plasticity and rapid adaptation to the environment. In particular, we show that global translation levels in CySCs depend on the self-renewal pathway JAK/STAT, and that CySCs and differentiated cyst cells have different requirements for translation initiation factors. The eIF4F complex is specifically required in CySCs for self-renewal but not differentiation, and acts genetically downstream of JAK/STAT signalling. We identify eIF3d1, the Drosophila homologue of eIF3d as a potential regulator of translation modes. We show that eIF3d1 is required for CySC self-renewal, like eIF4F, and that its phosphorylation by CkII is important for this function. Finally, we show that CkII acts downstream of JAK/STAT in CySCs, and that over-expressing CkII is sufficient to rescue both autonomous and non-autonomous roles of JAK/STAT signalling in CySCs. Altogether, we propose a model in which eIF3d1 acts as a switch between different modes of translation (Fig. 7): when JAK/STAT is active in CySCs, eIF3d1 is phosphorylated by CkII and enables eIF4F-dependent translation. We hypothesise that eIF4F drives selective translation of mRNAs encoding self-renewal factors, and does not translate mRNAs encoding differentiation factors, such as *eya.* As CySCs leave the niche and differentiate, JAK/STAT is inactive, leading to decreased CkII activity and consequently decreased eIF3d1 phosphorylation. Under these circumstances, eIF4F is not engaged in translation, and mRNAs encoding differentiation factors are translated.

**Figure 7.**
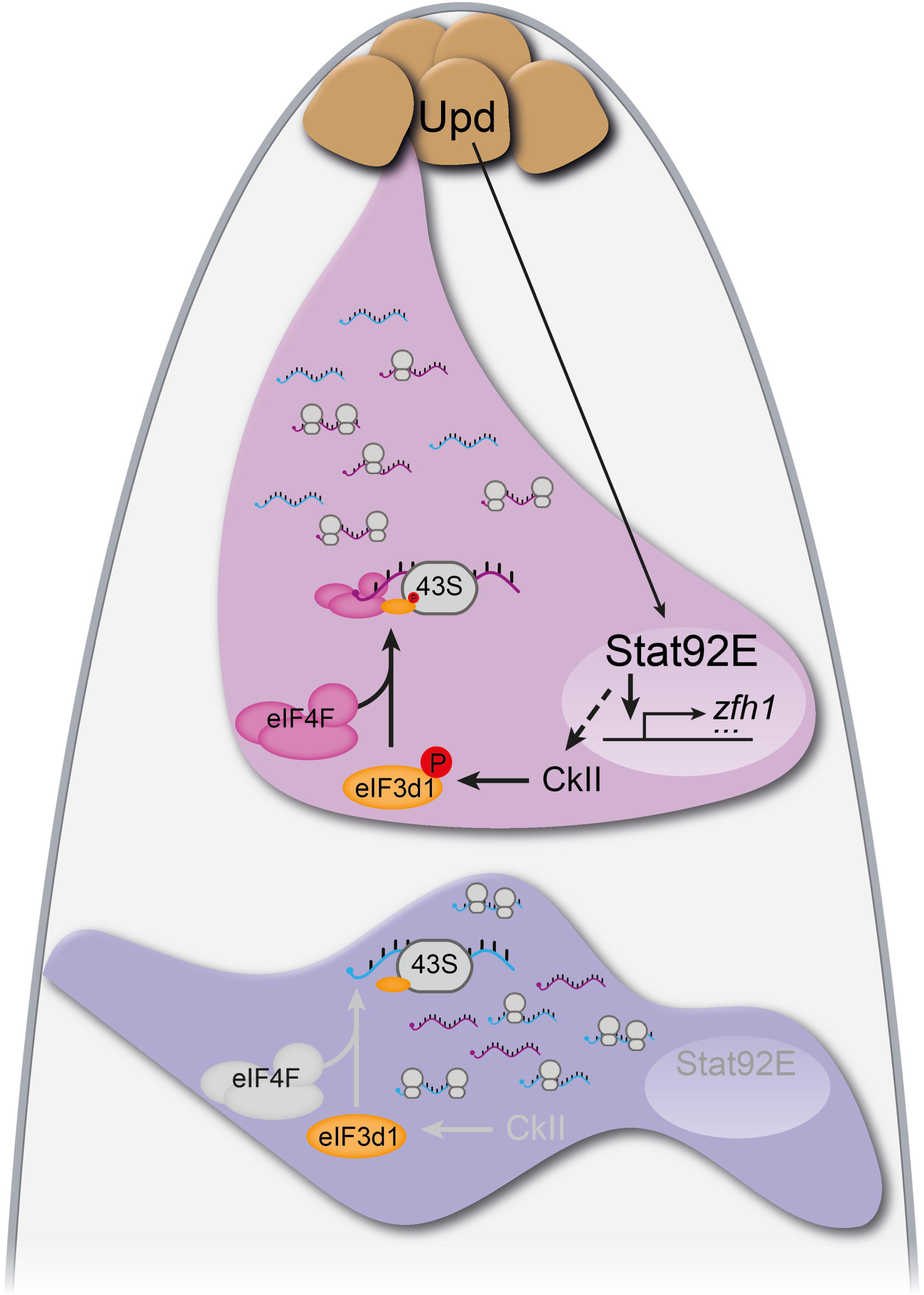
Model of translational control of stem cell fate in CySCs. The hub (brown) produces Upd which leads to the stabilisation and nuclear translocation of Stat92E in CySCs (magenta). Stat92E drives transcription of genes encoding self-renewal factors, including zfh1. Additionally, Stat92E promotes the activity of CkII, through targets which are currently unknown. In turn, CkII phosphorylates the translation initiation factor eIF3d1, which enables the interaction between eIF3d1 and eIF4F, leading to canonical cap-dependent translation. Our data suggest that eIF4F and eIF3d1 together preferentially translate mRNAs encoding self-renewal factors (purple), while mRNAs encoding differentiation factors (cyan), although present in CySCs, are not translated. In differentiating cells (blue), Stat92E is not found in the nucleus and CkII is inactive, such that eIF3d1 is unphosphorylated. In this situation, we propose that eIF3d1 no longer interacts with eIF4F, allowing cells to translate mRNAs independently of eIF4F. This results in changed specificity for the mRNAs that are bound to ribosomes, and preferential translation of differentiation factors instead of self-renewal factors.

### Different modes of translation initiation during stem cell differentiation

Our work indicates that stem cells and differentiated cells have different requirements for initiation factors. Cells lacking eIF4F appear to differentiate into functional cyst cells, suggesting it is dispensable and that translation initiation in cyst cells does not require canonical cap-binding activity. This is consistent with the global decrease in translation rates during cyst cell differentiation, as cap-independent translation mechanisms are thought to be less efficient than canonical initiation by recruitment of the ribosome through eIF4F. Intriguingly, although global translation rates change differently in different stem cell populations (for instance haematopoietic stem cells have lower translation rates than their differentiating offspring, while in Drosophila, both intestinal stem cells and CySCs have higher global translation), a common feature appears to be that translation is lower when cells finally become post-mitotic (Baser et al., 2019; Blanco et al., 2016; Obata et al., 2018; Signer et al., 2014; Wang and Amoyel, 2022). This suggests that the lower requirement for eIF4F activity may be conserved across differentiated cells in multiple tissues. Several studies have found that modulators of eIF4F activity, in particular 4E-BPs, can influence stem cell fate, although most have found that eIF4F activity is lower in stem cells (Hartman et al., 2013; Signer et al., 2016; Tahmasebi et al., 2014). While in some cases, this regulation has been linked to lower global translation, in ES cells, eIF4F regulation has little effect on global translation rates but affects specific transcripts (Tahmasebi et al., 2014; Tahmasebi et al., 2016). Intriguingly, unlike 4E-BP, Death Associated Protein 5 (DAP5, also called Novel APOBEC1 target 1 (Nat1) or eIF4G2), which interacts with eIF4A and eIF3, but lacks eIF4E binding and therefore cannot mediate cap-dependent translation, is required for differentiation of ES cells, both in mouse and humans, suggesting that translation of eIF4F-independent transcripts is required for differentiation (Sugiyama et al., 2017; Yoffe et al., 2016). Nonetheless, no study to date had evaluated the importance of different initiation factor complexes in stem cell fate determination, and our findings suggest that eIF4F may play specific roles in stem cell self-renewal, while it is dispensable during differentiation. Our measurements of OPP incorporation suggest that absolute levels of protein synthesis do not drive cell fate decisions (Fig. 2J,K). Thus, the importance of changing the initiation mode does not lie in the effect on global translation, rather, the mode of initiation must also determine the specificity of which mRNAs are translated. This provides buffering to the cell, ensuring noise in transcription does not influence cell fate. Indeed, we find transcripts encoding the differentiation factor *eya* present in CySCs when the protein is absent, suggesting that such a mechanism occurs in the Drosophila testis. In other tissues, there is extensive evidence from ribosomal profiling experiments that only a fraction of the transcriptome is translated (Baser et al., 2019; Ingolia et al., 2011; Lu et al., 2009; Schwanhausser et al., 2011; Spevak et al., 2020). Beyond just acting as a buffer for transcriptional noise, an intriguing hypothesis is that decoupling transcription and translation in this way could explain more generally the difference between specification and commitment during developmental trajectories. Specification is a state in which cells acquire a fate, but are still labile and can change fate if the environment instructs them to do so (Slack, 1991). How such plasticity is acquired is still extremely poorly understood; we speculate that overlaying a translational regulation programme onto transcriptional regulation of developmental gene expression could be a critical mechanism to allow cells to acquire a fate transcriptionally, but only commit to it when a second signal instructs a translational change that leads to a new proteome being synthesised and irreversible commitment to a particular differentiated fate. Of note, although specification is best understood in the context of early embryo development, there is evidence that adult stem cells are specified prior to differentiation, albeit under different names (“priming” or “licensing”), both in the Drosophila testis and in mammalian tissues (Nakagawa et al., 2021; Yuen et al., 2021). An important question is how different initiation factors provide specificity in translation. There is growing evidence that initiation factors are not just a passive machinery that translate all mRNAs, but that instead they have preferences either for the sequences around the cap, or for particular ternary structures in the mRNA (Leppek et al., 2018). Future work will determine the transcripts in CySCs that are present but not translated until cells begin differentiating to identify whether there are particular sequence or structural motifs that can determine the mode of initiation for a particular transcript. Additionally, although there are several mechanisms that are known to allow for translation in the absence of eIF4F (Kwan and Thompson, 2019), our data showing that knockdown of other initiation factors leads to ectopic self-renewal are harder to explain, given how crucial those factors are thought to be in all modes of translation initiation. One possibility is that by using RNAi, we only knocked down gene function incompletely. Another is that these disruptions lead to the activation of cellular stress pathways; there is evidence that activation of stress signalling can promote ectopic self-renewal (Biteau et al., 2008; Chang et al., 2019; Wang et al., 2014). Thus, unlike the clear effect of loss-of-function of eIF4F, it is still to be determined whether other translation factors drive specific gene expression programmes or whether they are simply required for all translation and lead to non-specific effects upon depletion.

### eIF3d enables a translational switch in response to environmental signals

Our work highlights the role of eIF3d1, the Drosophila homologue of eIF3d, as a regulatory switch controlling different translation modes. We show that phospho-dead eIF3d1, but not a form of eIF3d1 that lacks the mRNA 5’ cap-binding domain, acts as a dominant-negative for CySC self-renewal, and moreover, that only full length or phospho-mimetic eIF3d1 can rescue self-renewal upon knockdown of CkII. Given that eIF3d1 knockdown phenocopies knockdown of eIF4F components, these data suggest that eIF3d could mediate the interaction between the eIF3-containing 43S ribosome complex to mRNA-bound eIF4F, in a manner that depends on phosphorylation of eIF3d1. In this model, phosphorylation of eIF3d1 would act as a switch to selectively engage eIF4F-dependent translation. Indeed, our experiments using human cells support this hypothesis, showing that Ck2 inhibition weakens the interaction between eIF3d and eIF4F. Although in some situations, eIF3d has been shown to direct translation of specific transcripts (de la Parra et al., 2018; Lee et al., 2015; Lee et al., 2016; Volta et al., 2021), in the testis, its role appears to be similar to that of eIF4F, suggesting that it acts to maintain eIF4F activity, and that specific sequence recognition by eIF3d is less relevant to CySC fate decisions. Previous work has shown regulation of eIF3d through phosphorylation, demonstrating that upon glucose deprivation in cells, eIF3d is dephosphorylated and can bind the mRNA cap directly (Lamper et al., 2020). However, that work did not assess the interaction between eIF4F and eIF3d as it used a paradigm in which eIF4F activity was independently inhibited. In support of our model, studies have established that eIF3d and eIF4G can directly interact and that the presence of eIF3d is essential for the interaction between the core eIF3 octamer complex and eIF4G (Brito Querido et al., 2020; Sun et al., 2011; Villa et al., 2013). Alternatively, it has been suggested that the binding of eIF3d to internal structures within mRNAs could sterically inhibit eIF4F from associating with the 5’ cap (Ma et al., 2023). Therefore, it is possible that phosphorylation of eIF3d influences its ability to bind RNA instead of eIF4F, and that the effects on eIF4F activity are indirect. Determining how phosphorylation affects both canonical and non-canonical aspects of eIF3d activity will be important to understand how it can change the translational programme of cells.

Here we link the self-renewal signal from the niche and translational control via eIF3d1 and eIF4F. This provides an important link to explain how stem cells and differentiating progeny can have different levels of translation or specific translational programmes. While previous work has shown in several stem cell types that translation is selectively regulated, there has been little indication to date as to how signals from the niche could alter translation to influence fate. Here, we suggest a model that could apply to many other situations, and indeed beyond stem cell biology, to cancer, where translation, and increasingly eIF3d, have been shown to sustain cell proliferation and survival.

## Materials and methods

### Fly husbandry

Somatic-specific overexpression and knockdown experiments were carried out by using the *tj-Gal4; Tub>Gal80^ts^* system (referred to as *tj^ts^*) to allow temporal control of target gene expression in adult flies (McGuire et al., 2004). Crosses were raised at 18°C which is permissive for Gal80 activity. Males were collected 0-3 days after eclosion and shifted to the restrictive temperature of 29°C. Unless otherwise described in the text, flies were kept at 29°C for 10 days.

For clonal analyses, flies were raised at 25°C. Adult flies were collected 1-3 days after eclosion and heat shocked in a water bath at 37°C for 1 hour. Clonal CySCs were identified as labelled Zfh1-positive cells adjacent to the hub while clonal cyst cells were identified as either Zfh1-positive cells in the 2^nd^ row from the hub or Eya-positive cells.

For all experiments, *Stat92E^ts^* refers to the allelic combination *Stat92E^Frankenstein^/Stat92E^397^*. Recombinant chromosomes carrying *eIF4E* or *eIF3d1* mutations together with an FRT site were generated by crossing the relevant stocks and screening for resistance to neomycin and lethality.

### Generation of eIF3d overexpression flies

We used the Group-based Prediction System (GPS) server (Chen et al., 2023) to predict the phosphorylation sites in Drosophila eIF3d1, and compared to the human eIF3d phosphorylation sites reported in (Lamper et al., 2020) on Uniprot. DNA sequences encoding eIF3d^WT^, eIF3d^DD^ and eIF3d^NN^ were synthesized by Invitrogen GeneArt and cloned into pUAST-attB vector (RRID: DGRC_1419). Cloned plasmids were then injected by BestGene Inc into embryos from a strain carrying the *P{[+t7.7]=CaryP}Msp300[attP40]* landing site and inserted into the genome using PhiC31-mediated recombination.

### Immunohistochemistry

Dissected fly abdomens were fixed in 4% paraformaldehyde in PBS for 15 minutes then were washed twice in PBS, 0.5% Triton X-100 for 30 minutes for permeabilization. Permeabilized samples were blocked in PBS, 1% BSA, 0.2% Triton X-100 (PBTB) for one hour and incubated overnight in primary antibodies diluted in PBTB. Samples were washed twice in PBTB for 30 minutes each, and incubated in secondary antibodies diluted in PBTB for 2 hours at room temperature, followed by washes in PBS, 0.2% Triton X-100. Testes were separated from abdomens and mounted on slides with Vectashield mounting medium for imaging. We used the following antibodies: chicken anti-GFP (1:500, Aves Labs), rabbit anti-GFP (1:500, Invitrogen), rabbit anti-Stat92E (1:500, gift of E. Bach), guinea pig anti-Tj (1:3000, gift of D. Godt), rabbit anti-Zfh1 (1:5000, gift of R. Lehmann), guinea pig anti-Zfh1 (1:5000, this study). Mouse anti-Eya (eya10H6, 1:20, deposited by S. Benzer/N.M. Bonini), mouse anti-Fas3 (7G10, 1:20, deposited by C. Goodman), rat anti-CadN (1:20), rat anti-De-cad (1:20), rat anti-Vasa (1:20, deposited by A.C. Spradling/D. Williams) and mouse anti-Dlg (4F3, 1:20, deposited by C. Goodman) were obtained from the Developmental Studies Hybridoma Bank created by the NICHD of the NIH and maintained at The University of Iowa.

The guinea pig Zfh1 antibody was generated by GenScript. Recombinant antigen consisting of amino acids 648-775 of Zfh1 isoform PB was produced with an N-terminal His-tag used for purification. This antigen was injected into two guinea pigs. The resulting serum was purified by antigen affinity column to obtain concentrated antiserum.

For EdU or OPP staining, abdomens were dissected in Schneider’s medium instead of PBS and incubated for 30 minutes at room temperature in Schneider’s medium containing 10μM EdU or 5μM OPP. Samples were then fixed, permeabilized and stained with primary and secondary antibodies as above. Click reaction was carried out for 30 minutes at room temperature in the following reaction buffer: 2.5μM Alexa picolyl azide (Click Chemistry Tools), 0.1 mM THPTA, 2 mM sodium ascorbate and 1 mM CuSO4.

### In situ hybridization chain reaction (HCR)

HCR was carried out as previously described in (Duckhorn et al., 2022). 20 pairs of probes were designed, tiled along the *eya* transcripts and excluding regions of high similarity with other genes, with initiator sequences corresponding to amplifier B3 for amplification (Choi et al., 2018). Probes were purchased from ThermoFisher as DNA oligos, sequences are listed in File S1. Adult abdomens were dissected, fixed, and washed with PBS, 0.5% Triton X-100 as detailed above. Afterward, samples were incubated in Probe Hybridisation Buffer (Molecular Instruments) for 30 min at 37°C in water and incubated with pre-mixed probe pairs (0.01 µM for each probe) at 37°C overnight. Abdomens were then washed four times for 15 min each with Probe Wash Buffer at 37°C followed by a wash for 10 min with 5X saline sodium citrate solution: (SSCT): 14.61 g/mol sodium chloride, 73.53 g/mol 560 sodium citrate, pH 7, with 0.001% Tween 20 at room temperature. Samples were then incubated with Amplification Buffer for 10 min at room temperature. Meanwhile, 12 pmol of hairpins H1 and H2 were snap-cooled (heat at 95°C for 90 s and cooled to room temperature for 20 min) separately to prevent oligomerisation. The snap-cooled hairpins were then added to the samples in the Amplification Buffer and incubated overnight at room temperature. On the following day, samples were washed with 5X SSCT for 10 min then incubated with DAPI for 2h. After washing with ×1 PBS for 30 min, testes were mounted on slides as above.

### Cell culture and immunoprecipitations

2x10^6^ HeLa cells stably expressing Flag-tagged eIF3d were seeded in 10cm petri dish overnight. For the negative control, 1.5x10^6^ HeLa cells were seeded in 10 cm dish two days prior to the experiment, followed by transfection of Flag-RAP2A on the next day with the use of Lipofectamine 2000 (Thermo Fisher, #12566014). On the following day, cells were treated for 2 hours either with 3µM CX-5011 (AOB1816-5, Aobious) or DMSO, followed by quick wash with PBS and lysis with 400 µM of lysis buffer (50 mM Tris pH7.5, 150 mM NaCl, 1% Triton-X supplemented with 2x protease inhibitor cocktail (Roche, 11836145001), 1x phosphatase inhibitor cocktail (Roche, 11836170001), Sodium fluoride (50 mM), glycerol 2-phosphate (1 g/l), Sodium vanadate (2 mM), and Benzonase (50 U/ml)). Protein concentration of lysates was estimated with Pierce BCA (Life technologies, 23224, 23228). Based on the observed concentrations samples were balanced, a tenth of the samples was set aside and mixed with 5X Laemmli buffer to serve as the input control. The rest of the sample was mixed with anti-flag-beads (Sigma, A2220-5ml), which were prewashed three times with lysis buffer. The sample was incubated with the beads for two hours, rotating at 4°C. To remove unbound fraction, the sample was washed three times with 500 µl of lysis buffer.

Prior to the last wash the beads were moved into a new tube. Elution was performed by boiling the beads with 100 µl of Laemmli solution. The samples were further analysed by conventional western blot assay. We used the following antibodies: anti-eIF4G1 (Cell Signaling #8701), anti-eIF4A (Cell Signaling #2490), anti-eIF3b (Santa Cruz, discontinued), phospho-CK2 Substrate (pS/pT) (Cell Signaling #8738), anti-FLAG (Sigma, F7425), all at 1:1000.

## Statistical analysis

We used GraphPad Prism software to analyse data and generate graphs. For statistical analysis, we either used non-parametric ANOVA (Kruskal-Wallis test) followed by Dunn’s multiple comparisons test, to compare CySC numbers when there were several genotypes to compare, or Mann-Whitney tests for direct comparisons of CySC numbers when there were only two conditions. For normalized OPP level comparison, we used either parametric ANOVA (Šidák multiple comparisons test) or students’ t-test. Chi-square tests were applied for categorical data. Statistical tests used in each experiment are indicated in the relevant figure legends.

## Author contributions

Conceptualization, MA and RW; methodology, MA, RW, MR, AT; investigation, RW, MR, FS; writing – original draft, MA, RW; writing – review and editing, MA, RW, AT; supervision, MA, AT; funding acquisition, MA.

## Declaration of interests

The authors declare no competing interests.

## Supporting information

File S1 - probe sequences

## Acknowledgements

We thank S. Rumpf, E. Bach, T. Xie, R. Lehmann and D. Godt and the Bloomington, Vienna and Kyoto Drosophila stock centres for fly stocks and reagents. The authors are grateful to Ivana Bjedov, Nazif Alic, Richard Poole, Deepika Vasudevan, Vilaiwan Fernandes and members of the Amoyel and Fernandes labs for critical discussions and comments on the manuscript.

## Notes

### Competing Interest Statement

The authors have declared no competing interest.

