## Supplemental table and figures for "Niche signalling regulates eIF3d1 phosphorylation to promote distinct modes of translation initiation in stem and differentiating cells"

Supplementary information.

**Table S1 – Summary of initiation factor screen.**

Percent of testes with ectopic CySCs (assessed as Zfh1-positive, Eya-negative cells at least 3 cell diameters from the hub), or fewer than 10 CySCs near the hub. Note that some knockdowns with ectopic CySCs also occasionally led to a complete absence of somatic cells, so when ectopic cells were observed, the reduction of CySCs near the hub was not assessed.

| Knock-down | Stock number | % testes with ≤10 CySCs | Ectopic CySCs (%testes) | Number of testes examined |
|---|---|---|---|---|
| Control | NA | 0 | 0 | 26 |
| eIF4G1 | VDRC 17003 | 95 | 0 | 19 |
| eIF4A | VDRC 42202 | 100 | 0 | 17 |
| eIF4A | VDRC 100310 | 91 | 0 | 22 |
| eIF4E1 | VDRC 17581 | 27 | 0 | 22 |
| eIF4E1 | VDRC 7800 | 77 | 0 | 13 |
| eIF4E3 | VDRC 34210 | 0 | 0 | 21 |
| eIF4E4 | VDRC 107595 | 0 | 0 | 30 |
| eIF4E6 | VDRC 17580 | NA | 20 | 15 |
| eIF4H1 | VDRC 34301 | 0 | 0 | 19 |
| eIF4H2 | VDRC 102825 | 0 | 0 | 16 |
| eIF4B | VDRC 31364 | 0 | 0 | 15 |
| eIF3a | VDRC 28140 | NA | 80 | 15 |
| eIF3b | BDSC 32880 | NA | 86 | 14 |
| eIF3b | VDRC 107829 | NA | 82 | 31 |
| eIF3c | VDRC 26667 | NA | 40 | 5 |
| eIF3d1 | NIG 073-09 | 100 | 0 | 17 |
| eIF3d2 | VDRC 104342 | 0 | 0 | 16 |
| eIF3e | VDRC 27032 | NA | 41 | 22 |
| eIF3f | VDRC 101465 | NA | 50 | 22 |
| eIF3g | VDRC 28937 | NA | 65 | 23 |
| eIF3h | VDRC 106189 | NA | 78 | 37 |
| eIF3i | VDRC 27032 | NA | 90 | 20 |
| eIF2α | VDRC 7799 | NA | 75 | 40 |

| | | | | |
|---|---|---|---|---|
| eIF2γ | VDRC 39377 | NA | 60 | 15 |
| eIF1 | VDRC 29216 | NA | 16 | 18 |
| eIF1A | VDRC 26022 | NA | 41 | 22 |
| eIF6 | VDRC 108094 | NA | 35 | 17 |
| pAbp | VDRC 22007 | NA | 96 | 25 |
| eIF2D | BDSC 33995 | 0 | 0 | 16 |
| DENR | VDRC 28105 | NA | 5 | 18 |
| NAT1 | BDSC 27302 | 0 | 0 | 23 |

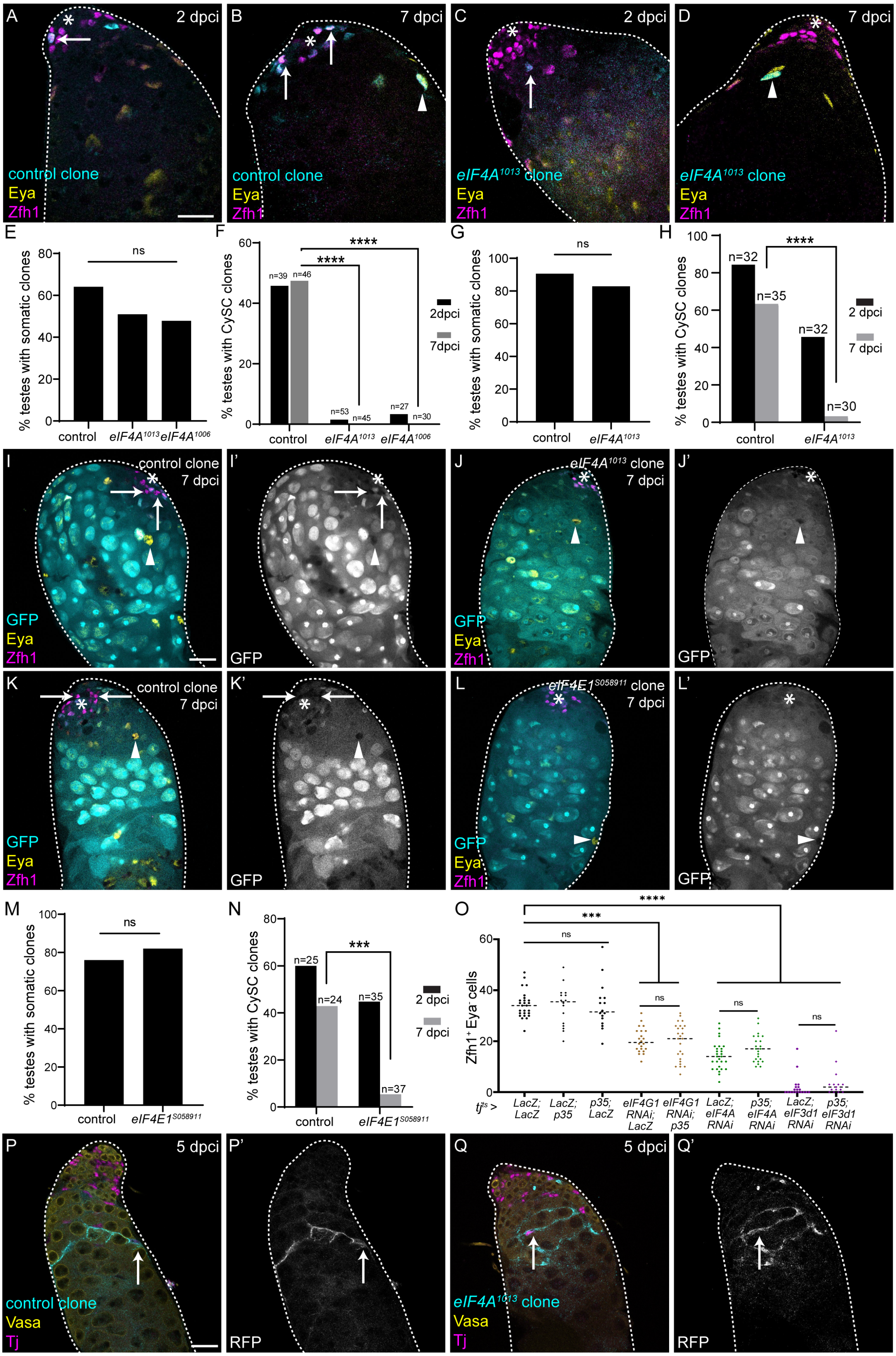

A 2 dpci control clone Eya Zfh1
B 7 dpci control clone Eya Zfh1
C 2 dpci eIF4A1013 clone Eya Zfh1
D 7 dpci eIF4A1013 clone Eya Zfh1
E % testes with somatic clones ns control eIF4A1013 eIF4A1006
F % testes with CySC clones **** **** n=39 n=46 n=53 n=45 n=27 n=30 2dpci 7dpci control eIF4A1013 eIF4A1006
G % testes with somatic clones ns control eIF4A1013
H % testes with CySC clones **** n=32 n=35 n=32 n=30 2 dpci 7 dpci control eIF4A1013
I control clone 7 dpci GFP Eya Zfh1
I' GFP
J eIF4A1013 clone 7 dpci GFP Eya Zfh1
J' GFP
K control clone 7 dpci GFP Eya Zfh1
K' GFP
L eIF4E1S058911 clone 7 dpci GFP Eya Zfh1
L' GFP
M % testes with somatic clones ns control eIF4E1S058911
N % testes with CySC clones *** n=25 n=24 n=35 n=37 2 dpci 7 dpci control eIF4E1S058911
O Zfh1+ Eya- cells **** *** ns ns ns ns tjts > LacZ; LacZ LacZ; p35 p35; LacZ eIF4G1 RNAi; LacZ eIF4G1 RNAi; p35 LacZ; eIF4A RNAi p35; eIF4A RNAi LacZ; eIF3d1 RNAi p35; eIF3d1 RNAi
P 5 dpci control clone Vasa Tj
P' RFP
Q 5 dpci eIF4A1013 clone Vasa Tj
Q' RFP

**Figure S1. eIF4F is specifically required for CySC self-renewal.**

(A-D) Testes with positively-marked control clones (A,B) or clones homozygous for *eIF4A*$^{1013}$ (C,D) at 2 dpci (A,C) and 7 dpci (B,D). GFP (cyan) labels the clone. Control clones are readily recovered in CySCs at 2 dpci and maintained at 7 dpci, while mutant clones are rarely observed adjacent to the hub. Zfh1 (magenta) labels CySCs and Eya (yellow) labels differentiated cells. Arrows mark CySCs and arrowheads mark differentiated cells. Asterisks indicate the hub. Scale bar: 15 μM.

(E) Percentage of testes with positively-marked control or *eIF4A* mutant clones in either CySCs or differentiated cyst cells at2 dpci (N≥27 testes, Chi-square test, ns P=0.3443).

(F) Percentage of testes with positively-marked control or *eIF4A* mutant CySC clones at 2 dpci and 7 dpci. (N≥27 testes, Chi-square test, **** P<0.0001).

(G) Percentage of testes with negatively-marked control or *eIF4A* mutant clones in either CySCs or differentiated cyst cells at 2 dpci (N≥30 testes, Chi-square test, ns P=0.3517).

(H) Percentage of testes with negatively-marked control or *eIF4A* mutant CySC clones at 2 dpci and 7 dpci. (N≥30 testes, Chi-square test, **** P<0.0001).

(I-J') Testes with negatively-marked control clones (E,E') and clones homozygous mutant for *eIF4A*$^{1013}$ (F,F') at 7 dpci. Clones are identified by lack of GFP (cyan). Mutant CySC clones are not recovered. Zfh1 (magenta) labels CySCs and Eya (yellow) labels differentiated cells. Arrows mark CySCs and arrowheads mark differentiated cells. Asterisks indicate the hub. Scale bar: 15 μM.

(K-L') Testes with negatively-marked control clones (G,G') and clones homozygous mutant for *eIF4E1*$^{S058911}$ (H,H') at 7 dpci. Clones are identified by lack of GFP (cyan). Mutant CySC clones are not recovered. Zfh1 (magenta) labels CySCs and Eya (yellow) labels differentiated cells. Arrows mark CySCs and arrowheads mark differentiated cells. Asterisks indicate the hub. Scale bar: 15 μM.

(M) Percentage of testes with negatively-marked control or *eIF4E1* mutant clones in either CySCs or differentiated cyst cells at 2 dpci (N≥24 testes, Chi-square test, ns P=0.3327).

(N) Percentage of testes with negatively-marked control or *eIF4E1* mutant CySC clones at 2 dpci and 7 dpci (N≥24 testes, Chi-square test, *** P<0.001).

(O) Number of Zfh1$^{+}$ Eya$^{-}$ CySCs in the testis upon knockdown of initiation factors and inhibition of apoptosis using the baculovirus caspase inhibitor P35. Blocking cell death did not rescue CySC loss caused by knockdown of initiation factors (N≥15 testes, Kruskal-Wallis test, **** P<0.0001, *** P<0.001 ns P>0.9999).

(P-Q') Testes with membrane-labelled control clones (P,P') and clones homozygous mutant for *eIF4A*$^{1013}$ (Q,Q') at 5dpci. RFP (cyan) labels the clones (arrows). Both control and mutant clones display the characteristic morphology of cyst cells, with a long flattened cytoplasm enveloping germ cell cysts (arrows). Tj (magenta) labels cyst cells and Vasa (yellow) labels germ cells. Scale bar: 15 μM.

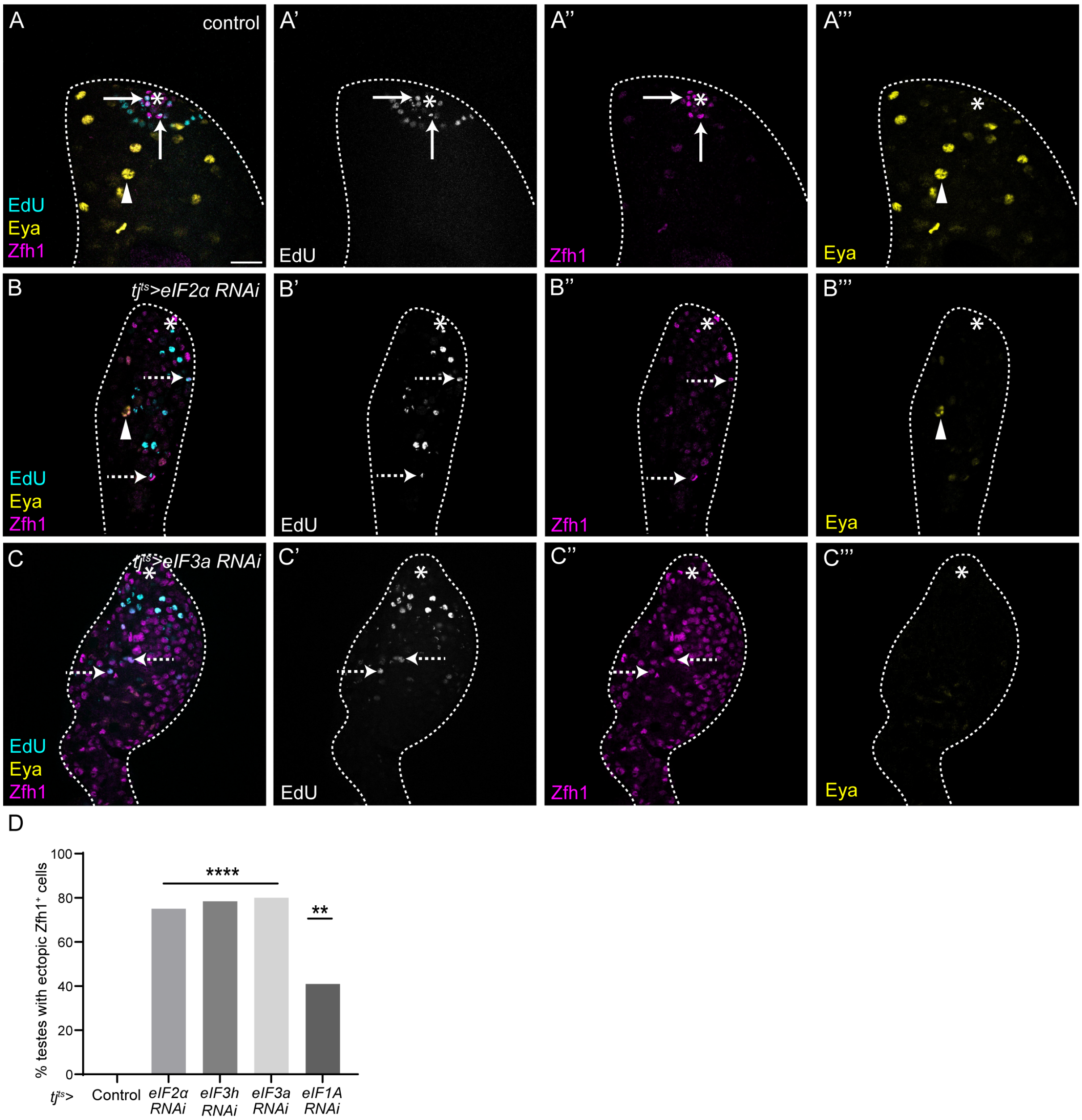


**Figure S2. Knockdown of initiation factors results in ectopic CySC-like cells.**

(A-C''') A control ($tj^{ts}$ > +) testis (A-A'''), and testes in which eIF2α (B-B''') or eIF3a was knocked down in CySCs (C-C'''). EdU (cyan) labels cells in S phase. In controls, EdU-positive somatic cells are found only adjacent to the hub, while in eIF knockdowns, EdU-positive CySC-like cells are observed away from the hub. Zfh1 (magenta) labels CySCs and Eya (yellow) labels differentiated cyst cells. Arrows mark CySCs, dashed arrows indicate ectopic Zfh1-positive cells away from the hub, and arrowheads mark differentiated cells. Asterisks indicate the hub. Scale bar: 15 µM.

(D) Percentage of testes with $Edu^+$ $Zfh1^+$ cells at least 2 cell diameters from the hub (N≥15 testes, Chi-square test, **** P<0.0001, ** P<0.01).

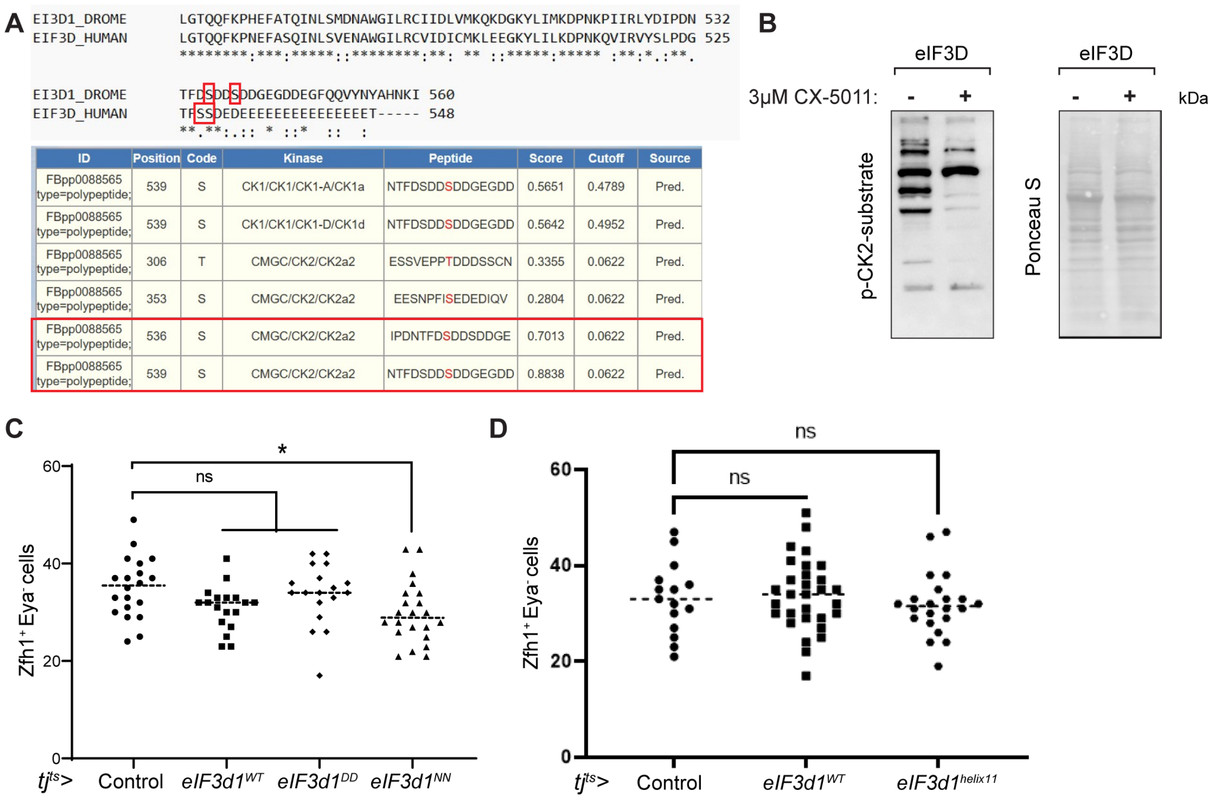


**Figure S3. eIF3d1 phosphorylation but not cap-binding is required for CySC self-renewal.**

(A) Top: alignment of the N terminus of human eIF3d and Drosophila eIF3d1. Asterisks indicate conserved residues, colons indicate residues with strongly similar properties and periods indicate residues with weakly similar properties. The two serines subject to phosphorylation in human eIF3d are highlighted in red boxes along with predicted phosphorylation sites in Drosophila. Bottom: predicted phosphorylation sites in Drosophila eIF3d1 using Group-based Prediction System (GPS) server. The boxed lines indicate serines 536 and 539, highlighted above, which are strong predicted targets of CkII.

(B) Western blot from lysates of cells expressing a control construct (RAP2A) or eIF3d and labelled with an antibody against a pan-phospho-CK2 substrate (left) or stained for total protein (right). Addition of the CK2 inhibitor CX-5011 resulted in decreased labelling for the phospho-CK2 substrate.

(C) Number of Zfh1$^{+}$ Eya$^{-}$ CySCs in the testis upon over-expression of wild type (eIF3d1$^{WT}$), phospho-mimetic (eIF3d1$^{DD}$) or phospho-dead (eIF3d1$^{NN}$) eIF3d1. Phospho-dead eIF3d1 expression results in a significant decrease in CySC numbers (N≥18 testes, * $P<0.05$, ns $P=0.1150$ (eIF3d1$^{WT}$) or $P>0.9999$ (eIF3d1$^{DD}$), Kruskal-Wallis test).

(D) Number of Zfh1$^{+}$ Eya$^{-}$ CySCs in the testis upon over-expression of wild type eIF3d1 or a form of eIF3d1 lacking the mRNA 5’ m$^{7}$G cap-binding domain (eIF3d1$^{helix11}$) (N≥15 testes, Kruskal-Wallis test, ns $P>0.9999$).

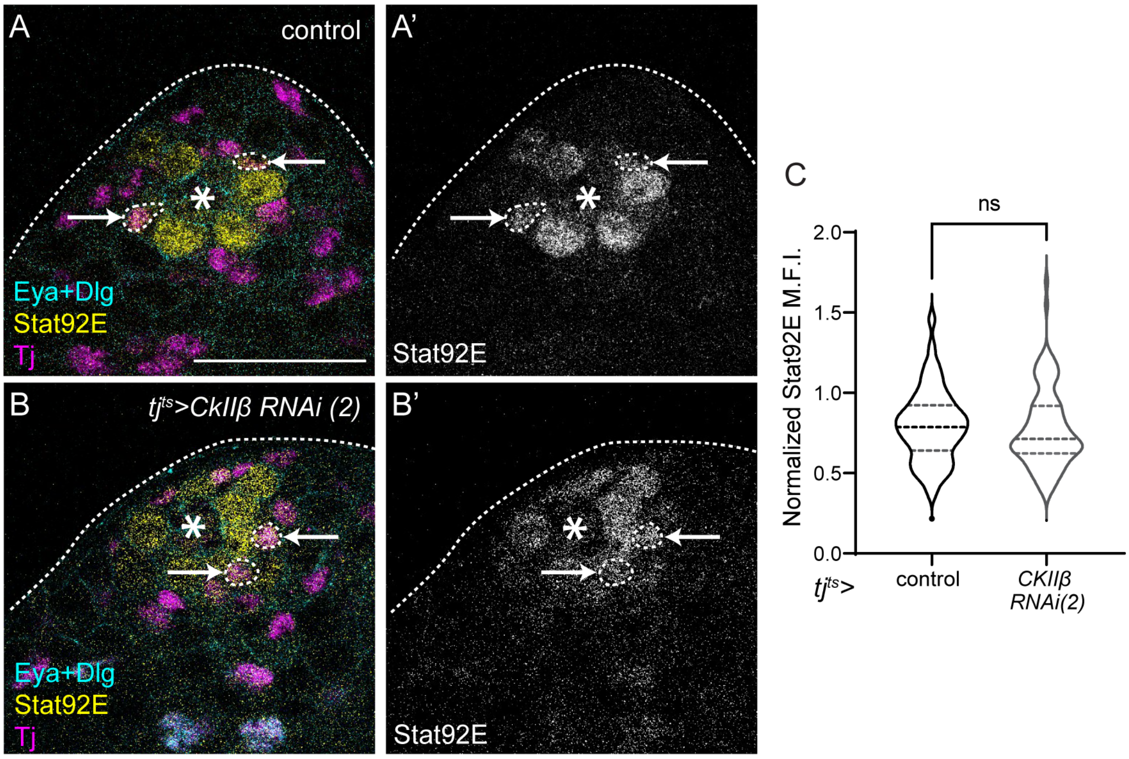


**Figure S4. CkII knockdown does not affect JAK/STAT signal transduction.**

(A-B) Control (A) and testis in which CkIIβ was knocked down in CySCs with *tj$^{ts}$-Gal4* (B), stained with an antibody against Stat92e (yellow, single channel A',B') after 20h at the restrictive temperature. Tj (magenta) labels CySCs and early cyst cells, Dlg (cyan) labels the hub and Eya (cyan) labels differentiated cyst cells. Arrows mark CySCs. Asterisks indicate the hub. Scale bar: 15 μM.

(C) Quantification of Stat92E levels in CySCs in control testes or upon CkIIβ knockdown, normalized to levels in neighbouring GSCs. (N□100 cells from □7 testes, Student's t-test, ns P=0.5057)
